# Mapping of corticotropin-releasing factor, receptors and binding protein mRNA in the chicken telencephalon throughout development

**DOI:** 10.1101/2023.03.07.531566

**Authors:** Alek H. Metwalli, Alessandra Pross, Ester Desfilis, Antonio Abellán, Loreta Medina

**Affiliations:** Dept. of Experimental Medicine, Universitat de Lleida, Lleida, Spain; Laboratory of Evolutionary and Developmental Neurobiology, Lleida’s Institute for Biomedical Research-Dr. Pifarré Foundation (IRBLleida), Lleida, Catalonia, Spain; Serra Húnter fellow

**Keywords:** hippocampus, amygdala, auditory pallium, visual pallium, hypothalamo-pituitary-adrenal axis, stress

## Abstract

Understanding the neural mechanisms that regulate the stress response is critical to know how animals adapt to a changing world and is one of the key factors to be considered for improving animal welfare. Corticotropin releasing factor (CRF) is crucial for regulating physiological and endocrine responses, triggering the activation of the sympathetic nervous system and the hypothalamo – pituitary – adrenal axis (HPA) during stress. In mammals, several telencephalic areas, such as the amygdala and the hippocampus, regulate the autonomic system and the HPA responses. These centers include subpopulations of CRF containing neurons that, by way of CRF receptors, play modulatory roles in the emotional and cognitive aspects of stress. CRF binding protein also plays a role, buffering extracellular CRF and regulating its availability. CRF role in activation of the HPA is evolutionary conserved in vertebrates, highlighting the relevance of this system to help animals cope with adversity. However, knowledge on CRF systems in the avian telencephalon is very limited, and no information exists on detailed expression of CRF receptors and binding protein. Knowing that the stress response changes with age, with important variations during the first week posthatching, the aim of this study was to analyze mRNA expression of CRF, CRF receptors 1 and 2, and CRF binding protein in chicken telencephalon throughout embryonic and early posthatching development, using in situ hybridization. Our results demonstrate an early expression of CRF and its receptors in pallial areas regulating sensory processing, sensorimotor integration and cognition, and a late expression in subpallial areas regulating the stress response. However, CRF buffering system develops earlier in the subpallium than in the pallium. These results help to understand the mechanisms underlying the negative effects of noise and light during prehatching stages in chicken, and suggest that stress regulation becomes more sophisticated with age.

## Introduction

Corticotropin-releasing factor (CRF) or hormone (CRH) and its receptors play critical roles in the regulation of fear and anxiety responses, and dysregulation of CRF systems are involved in several stress- and mood-related disorders (reviewed by **Steckler and Holsboer, 1999; Bale and Vale, 2004**; **Henckens et al., 2016**; **Dedic et al., 2018**; **Deussing and Chen, 2018**). During the stress response, this neuropeptide triggers the activation of the autonomic sympathetic nervous system and the hypothalamo-pituitary-adrenal axis (HPA) (**Bale and Vale, 2004**; **Henckens et al., 2016**; **Herman et al., 2016**); the first leads to liberation of adrenaline, and the second to the release of cortisol/corticosterone to blood, which acts on multiple organs to redirect energy resources and adapt to changes in the environment, while maintaining homeostasis in the organism (**Bale and Vale, 2004; Herman et al., 2016; Dedic et al., 2018**). HPA activation can occur during aversive negative situations, but also in response to an appetitive, rewarding stimulus (**Dedic et al., 2018**). Activation starts with the stimulation of CRF-expressing parvocellular neurons of the paraventricular hypothalamic nucleus (PVN), which release this factor into the portal venous system at the median eminence, thus reaching the anterior pituitary gland where it binds to CRF receptor 1 (CRFR1) and produces the secretion of adrenocorticotropin hormone (ACTH) to the blood stream; the latter acts on cortical cells of the adrenal glands, which release corticosterone to the blood system (**Bale and Vale, 2004**). Although CRF is the major activator of HPA (**Dedic et al., 2018**), vasopressin is co-released with CRF from terminals at the median eminence, and acts synergistically with it to increase the secretion of ACTH by pituitary corticotropes, an action that is mediated by vasopressin 1b receptors (**Bale and Vale, 2004; Herman et al., 2016**). The production of corticosterone is subjected to a complex regulation by the brain, which is affected by circadian rhythms, and by the levels of several substances in blood including inflammatory factors and gonadal steroids (**Herman et al., 2016**). Moreover, corticosterone regulates its own production and acts on the brain, through glucocorticoid/mineralocorticoid receptors, after passing the brain-blood barrier, giving rise to a potent negative-feedback that deactivates the HPA (**Bale and Vale, 2004**). Brain regulation of HPA is extremely intricate, involving several centers in the brainstem, hypothalamus, and telencephalon, as well as a variety of CRF and non-CRF neuron subpopulations and pathways (**Herman et al., 2016; Henckens et al., 2016**).

In mammals, many of the brain areas involved in stress and in regulation of HPA and/or the autonomic nervous system include subpopulations of CRF neurons, such as the prefrontal cortex, the hippocampus, the extended amygdala, and the preoptic area (**Swanson et al., 1983**; **Merchenthaler, 1984; Morin et al., 1999; Henckens et al., 2016**). In these areas, CRF appears to play modulatory roles in relation to arousal, feeding behavior, emotion, memory formation, and other cognitive aspects of stress (**Dedic et al., 2018**). The cell composition of these and other brain areas is heterogeneous, making it difficult to discern the specific role of each CRF cell type in stress response and their relationship to other neurons of the same area/nucleus. For example, both the central amygdala (in particular, its lateral subdivision) and the BSTL contain many different neuron types, including subsets of GABAergic neurons coexpressing CRF (**Gray and Magnuson, 1992**; **Marchant et al., 2007**; **McCullough et al., 2018**; **Kovner et al., 2019**). Interestingly, CRF expression and the activity of CRF cells in the central amygdala and BST is enhanced upon exposure to high levels of glucocorticoids and some chronic stress situations, such as chronic social subordination and chronic pain (**Herman et al., 2016; Partridge et al., 2016**).

In the brain, CRF acts through two types of G-protein coupled receptors, named CRFR1 and CRFR2. These receptors share 70% of their amino acid sequence, and variations mostly refer to their ligand-binding region (**Bale and Vale, 2004; Henckens et al., 2016**). CRF shows highest affinity for CRFR1, but both receptors also bind different urocortins, which belong to the CRF family and share 26 to 43% amino acid identity with CRF (**Dedic et al., 2018**). Studies in rodents have shown that signaling through these receptors regulates anxiety-like behavior in a region, cell type and circuit specific mode (**Henckens et al., 2016**). CRF effects also depend on stressor modality, dose and time of exposure, and age (**Chen et al., 2004a, 2006, 2010**; **Herman et al., 2016**; **Vandael et al., 2021**). These receptors show different expression patterns in the brain, with CRFR1 having a more widespread (but not ubiquitous) expression than CRFR2 (**Henckens et al., 2016**). Regarding CRFR1, in addition to the anterior pituitary, it shows moderate to high expression in several areas of the telencephalon (including neocortex, hippocampus, amygdala, and olfactory bulb), parts of hypothalamus (like the paraventricular nucleus), reticular prethalamic nucleus, cerebellum and numerous brainstem centers (**Chalmers et al., 1995; Chen et al., 2000**, **2004a**; **Henckens et al., 2016; Weera et al., 2022**). In contrast, CRFR2 shows high expression in the lateral septum, ventromedial hypothalamus, and choroid plexus, while its expression is moderate in the hippocampus, extended amygdala, paraventricular and supraoptic hypothalamic nuclei, midbrain colliculi and raphe nuclei (**Chalmers et al., 1995**; **Henckens et al., 2016; Dedic et al., 2018**). CRFR2 is not expressed in the cerebral cortex and anterior pituitary in rodents, but in humans this receptor is found in these locations (**Dedic et al., 2018**), pointing to variations between species that need to be considered to better understand the regulation of the stress response in different animals. CRF receptors are expressed in different cell populations that contain different classical neurotransmitters, including glutamatergic neurons (most of those in the telencephalic pallium and the paraventricular and ventromedial hypothalamic nuclei), GABAergic neurons (most of those in the telencephalic subpallium and the reticular prethalamic nucleus), serotonergic neurons (in the raphe nuclei), dopaminergic neurons (in the ventral tegmental area), noradrenergic neurons (in locus coeruleus), and cholinergic neurons (in the laterodorsal tegmental nuclei) (**Henckens et al., 2016**). Determining the specific function of different CRF receptors in distinct neuron populations and circuits has been technically difficult due to the heterogeneity of the areas, but this has improved in recent years thanks to the generation of reporter mice and rats, which combined with conditional mutagenesis, viral manipulation and optogenetics is allowing site- and cell-specific manipulation of different receptors to study their function in specific neuron populations and circuits (see details in **Henckens et al., 2016; Dedic et al., 2018; Weera et al., 2022**). These studies also need to consider the role of CRF binding protein, which binds to extracellular CRF and appears to buffer and regulate the availability of this peptide, thus modulating CRF function during stress related responses (**Bale and Vale, 2004; Dedic et al., 2018**).

Activation of HPA during the stress response follows similar rules in different vertebrates, including birds (**Carsia et al., 1986**; **Vandenborne et al., 2005**; **Smulders, 2021**), and this appears designed to help animals cope with adversity (**Herman et al., 2016**). In birds, it also involves activation of CRF neurons of the paraventricular hypothalamic nucleus, leading to release of ACTH by the pituitary and subsequent release of corticosterone by adrenal glands (**Carsia et al., 1986; Vandenborne et al., 2005**). The avian paraventricular hypothalamus also receives input from the extended amygdala, including the BST, where GABAergic neurons concentrate, and the latter receives input from the hippocampal formation (**Atoji et al., 2006**), suggesting the existence of a similar regulation of HPA by the telencephalon in birds.

Moreover, CRF cell distribution and fiber systems in the brain also appear to be similar between birds and mammals (**Józsa et al., 1984; Richard et al., 2004**). Chicken CRF peptide is identical to human and rat CRF (**Vandenborne et al., 2005**), and the gene that encodes it is orthologous to CRF1 of mammals and other vertebrates (**Cardoso et al., 2016, 2020**). In addition, like in other vertebrates (except teleost fishes and placental mammals), a second type of CRF (CRF2) is found in chicken, which is expressed in the brain (as determined by qPCR) and, at least in vitro, is able to bind to CRF receptors as follows: it shows high preference for and is a potent activator of CRFR2, and it has low affinity for CRFR1, being able to activate the latter only at high doses (**Bu et al., 2019**).

Like CRF peptides, CRF receptors (1 and 2) and binding protein (CRFBP) also are highly conserved in evolution (**Yu et al.., 1996**; **De Groef et al., 2004**; **Lovejoy and de Lannoy, 2013**; **Cardoso et al., 2016, 2020**; **Wan et al., 2022**). Using in situ hybridization in chicken, CRFR1 was specifically found in ACTH producing pituitary cells, while CRFR2 was expressed in TSH producing cells (**De Groef et al., 2003).** Studies by way of qPCR showed that mRNAs of both receptors are expressed in several brain regions of chicken, such as telencephalon, hypothalamus, midbrain tectum, and hindbrain (**De Groef et al., 2004**), while CRFBP mRNA is abundant in the telencephalon and hypothalamus, poor in the midbrain, and poor to moderate in the hindbrain (**Wan et al., 2022**). However, information on the detailed expression of CRF receptors and binding protein in the central nervous system is missing in birds, and thus many aspects on the degree of conservation of the brain mechanisms regulating the stress response are still unknown. This information is crucial to understand how stress is regulated in different animals, and to improve animal welfare. The aim of this study is to carry out a detailed mapping of mRNA expression of CRF receptors and binding protein in chicken telencephalon, compared to distribution of CRF expressing cells. Since the stress response changes with age in different animals (**Herman et al., 2016**), including birds, passing through a hyporesponsive period around birth or hatching (**Schapiro et al., 1962**; **Freeman and Manning, 1984**), we will analyze the dynamic expression patterns of CRF related peptides and receptors throughout embryonic and early posthatching development. The results can help to understand these transitions in stress responsivity, as well as the long-term negative effects of early-life stress (**Janczak et al., 2006**; **De Haas et al., 2021**).

## Material and Methods

### 1. Tissue collection, fixation and sectioning

Fertilized chicken embryos of the *Gallus gallus domesticus* (Leghorn) were obtained from a commercial hatchery (Granja Santa Isabel, Cordoba, Spain; Authorization ES140210000002), and then incubated in a draft-free incubator at 37.5 °C and 55-60% humidity until the desired stage. All animals were treated according to the regulations and laws of the European Union (Directive 2010/63/EU) and the Spanish Government (Royal Decrees 53/2013 and 118/2021) for the care and handling of animals in research. The protocols used were approved by the Committees of Ethics for Animal Experimentation and Biosecurity of the University of Lleida (reference no. CEEA 08-02/19), as well as that of the Catalonian Government (reference no. CEA/9960_MR1/P3/1 for embryos, and CEA/9960_MR1/P4/1 for post-hatchlings).

We used 28 embryos between 8 (E8; equivalent to Hamburger and Hamilton’s stages HH34/35) and 18 (E18, equivalent to HH44) days of incubation (E8, N=4; E12, N=2; E14, N=8; E16, N=3; and E18, N=11), plus seven neohatched (P0) and four 7 days-old (P7) posthatchlings, of both sexes. Younger embryos were rapidly decapitated and their brain dissected and fixed by immersion in 0.1 M phosphate buffered 4% paraformaldehyde (PFA). Older embryos (E14, E16, E18) and post-hatchlings were first anaesthetized with MS222 (tricaine methanesulfonate) or halothane (**Pross et al., 2022**), followed by sodium pentobarbital (with a euthanasic dose of 100 mg/kg, intraperitoneal), and then they were perfused transcardially with saline followed by PFA. After dissection, the brains were postfixed in PFA at 4°C, for 24h. Subsequently, the brains were washed 3 times with 0.1 M phosphate buffered saline (PBS, pH 7.4) for 10 min, under mild shaking, and cryopreserved at -20°C in hybridization buffer before further use.

Immediately before being used, brains were embedded in a 4% low-melt agarose matrix and sectioned in frontal, sagittal and horizontal planes using a Leica VT 1000S vibratome (speed 4-5, thickness 100 µm). Sections were serially collected in PBS, and kept at 4°C. Sections were processed for in situ hybridization to study expression of chicken genes encoding corticotropin-releasing factor (Crf), Crf receptors 1 and 2 (Crfr1, Crfr2), and Crf binding protein (Crfbp).

### 2. Preparation of the riboprobes for *in situ* hybridization experiments

The clones were obtained by BBSRC ChickEST Database (Boardman et al., 2002) and distributed by Source BioScience, except for that of Crf, which was bought from GenScript (Table 1).

**Table 1:** Genes and cDNAs employed for in situ hybridization

| **Gene name** | **Gene accession number** | **EST (expressed sequenced tag) code** | **Size (bp) and alignment with accession number** | **Obtained from** |
| --- | --- | --- | --- | --- |
| Chicken Corticotropin releasing factor (*Crf*) | NM_001123031.1 |  | 504 (165-668) | GenScript |
| Chicken Corticotropin releasing factor receptor 1 (*Crf1*) | NM_204321 | ChEST704a13 | 672 (899-1571) | Source BioScience |
| Chicken Corticotropin releasing factor receptor 2 Variant X2 (*Crf2*) | XM_015281045.4 | ChEST43i8 | 899 (1674-2387) | Source BioScience |
| Chicken Corticotropin releasing factor binding protein Variant X2 (*Crfbp*) | 5XM_046935644.1 | ChEST66a | 858 (14-858) | Source BioScience |

#### 2.1. Plasmid production, linearization, amplification and purification

The clones were used to transform bacteria following the protocol of Invitrogen for DH5α competent bacterial (ref. 18265-017; Subcloning Efficiency™ DH5α Competent Cells; Invitrogen ThermoFisher Scientific). The bacteria containing the plasmid of interest (Table 1) were grown overnight in 50 ml of liquid LB culture medium (1% Bacto-triptone; 0.5% yeast extract; 1% NaCl and 10 mg/ml of ampicillin). After, the plasmids were isolated using the Qiagen Plasmid Midi Kit (ref. 12243).

Plasmids obtained in the previous step were linearized and/or amplified following two methods, as follows. Those containing the insert of Crf were directly amplified using PCR. The rest of the clones were linearized using restriction enzymes bought from Takara, using a standard procedure. This was followed by amplification by PCR, using *Taq* PCR Master Mix (Qiagen; ref 201445) and two primers containing the sequence of T7 (5’-CGTAATACGACTCACTATAGGGCGA-3’) and T3 ((5’-GCAATTAACCCTCACTAAAGGGAAC-3’) promotors, present in the pBlueScript II SK (+) plasmid.

Crf containing clone was acquired from GenScript, but the vector (pcDNA3.1-C-(k)DYK) did not present the T3 promotor to synthetize antisense RNA, but only the T7 for sense. This problem was solved by amplifying by PCR the insert with a new primer containing the T3 promotor site, allowing to synthesize antisense RNA. For it, we used the Phire Hot Start DNA polymerase (Finnzymes-Thermo Fisher Scientific; ref. F-120) and the primer 5’-GATATTAACCCTCACTAAAGGGAAC CTTCCGATGATTTCCATCAGTTTC-3’, which contains a piece of the Crf gene sequence and the T3 promotor sequence to polymerize antisense RNA. The other primer contained the T7 binding sequence (as explained above). The size differences between both primers were solved using two annealing temperatures (51 °C for 10 cycles and 56°C for 20 cycles). Linearized DNAs obtained from the previous steps were purified following the instructions of UltraClean PCR Clean-Up Kit (MO BIO Laboratories; ref 12500-100).

#### 2.2. Synthesis and purification of the riboprobe

Linearized DNAs were used as templates to synthesize *in vitro* antisense RNA following the protocol of the RNA polymerase protocol (Roche Diagnostic). The mix containing the DNA, digoxigenin-labeled nucleotide mix and the specific RNA polymerase was incubated for 2h at 37°C, and later it was purified by adding 100 µl TE buffer (10mM Tris-HCl pH8; 0,1mM ethylenediaminetetraacetic acid (EDTA), 5 µl 8M LiCl (Sigma-Aldrich) and 300 µl of absolute ethanol molecular grade pure (Scharlau). The resulting mix was placed at -20°C for at least 30 minutes before proceeding with the purification of the probe.

The mix obtained in the previous step was centrifuged for 30 minutes at 14000g at 4°C, then the supernatant was discarded, the pellet was washed with 200 µl of 70% Ethanol (Scharlau), followed by centrifugation for 10 minutes at 14000g at 4°C. The supernatant was discarded and the pellet was left to dry for 2-5 minutes. Thereafter, the pellet was resuspended in the 25 µl of molecular water (DNAase and RNAase free; Sigma-Aldrich), and, finally, conserved by adding 25 µl of molecular formamide to avoid solidification when stored at -20 °C.

### 3. *In situ* hybridization

**First Day**. Free floating sections were prehybridized in the prehybridization buffer for 2 hours at 58°C and then hybridized in the hybridization buffer, containing the riboprobe overnight at 63°C (0,5-1µg/ml depending on the probe). The composition of buffers was as follows:

Prehybridization buffer: 50% formamide for molecular use (Fisher); 1.3× standard saline citrate (pH 5); 5 mM ethylenediaminetetraacetic acid (EDTA pH 8.0; Sigma-Aldrich), 1 mg/ml of yeast tRNA (Sigma-Aldrich); 0.2% Tween-20 and 100 μg/ml of heparin (Sigma-Aldrich), completed with molecular water (free of RNAase and DNAase; Sigma-Aldrich).

Hybridization buffer: 50% formamide molecular (Fisher); 10% Dextran Sulfate (Sigma); 1 mg/ml of yeast tRNA (Sigma-Aldrich); 0.2% Tween-20; 1x Denhardt Solution (Sigma-Aldrich); 1 x Salt Solution (Sigma-Aldrich), completed with molecular water (free of RNAase and DNAase; Sigma-Aldrich).

**Second day**. After hybridization, the sections were washed abundantly first with hybridization buffer of second day (HB2) at 63°C during 30 minutes and, then, 30 minutes at room temperature (RT) under mild shaking. HB2 contained 50% formamide (Sigma-Aldrich); 1.3X standard saline citrate (pH 5); 5 mM ethylenediaminetetraacetic acid (pH 8.0; Sigma-Aldrich); 1 mg/ml of yeast tRNA (Sigma-Aldrich); 0.2% Tween-20, in water (free of RNAase and DNAase; Sigma-Aldrich). The second wash was with 1:1 mix of MABT (1.2% maleic acid, 0.8% NaOH, 0.84% NaCl and 0.1% Tween-20) and BH2 at 58°C for 20 min, followed by 15 min at room temperature under mild shaking. After this, the tissue was washed for 2 hours with MABT, changing the buffer every 30 minutes, at room temperature. Then, sections were blocked in a saturation solution containing 2% Blocking Reagent (Roche Diagnostics) and 20% sheep serum in MABT for 4h at room temperature, and then incubated in an anti-digoxigenin antibody (alkaline-phosphatase-coupled anti-digoxigenin; RRID:AB_514497, diluted 1:3,500; Roche Diagnostics; Table 2) for 30 minutes at room temperature, followed by overnight incubation at 4°C.

**Table 2:** Antibody

| Primary antibody |  |  |  |  |
| --- | --- | --- | --- | --- |
| Antibody name | Type | Antigen recognized | Dilution | Manufacturer and reference |
| Anti-Digoxigenin-AP Fab fragments | Polyclonal | Digoxigenin | 1:3500 | Roche, ref 11093274910<br>RRID:AB_514497 |

**Third day**. The antibody solution was removed, and sections were washed with 0.1M PBS containing 0,3% Triton-100 (PBS-T) during 2 hours at room temperature, under mild shaking, changing the washing solution every 30 minutes. Then, sections were washed two times with MABT and B2 buffer (100mM Tris-HCl, 100mM NaCl, 50mM MgCl2, pH 7.5) for 30 minutes each one. The hybridization signal was revealed with NBT (nitroblue tetrazolium) and BCIP (5-bromo-4-chloro-3-indolyl phosphate) (both from Roche Diagnostics) mixed in B2 in a proportion of 0.45 µl of NBT and 4.5 µl of BCIP in 5 ml of B2. To finish the development, the sections were washed with MABT and we stopped the reaction with 4% paraformaldehyde.

Finally, the sections were mounted on glass slides with 0.5% Gelatine from porcine skin (Sigma-Aldrich) in 0.1 M Tris-Cl buffer (pH 8), and when the sections were dry, they were cover slipped with either Glycerol Gelatin (Sigma-Aldrich) without dehydration, or Permount (Fisher Scientific) previous dehydration with alcohols.

### 4. Digital photographs and figures

Digital microphotographs from hybridized sections were taken on a Leica microscope (DMR HC, Leica Microsystems GmBH) equipped with a Zeiss Axiovision Digital Camera (Carl Zeiss, Germany). The figures were mounted with selected images by means of CorelDraw 2019 (Corel Corporation, Canada).

### 5. Nomenclature

For the identification of forebrain cell masses, we primarily followed the proposal of the Avian Brain Nomenclature Forum (**Reiner et al., 2004**) and the chick brain atlas (**Puelles et al., 2019**). For the developing chicken brain, we followed **Puelles et al. (2000**) as well as our own publications on the subject (**Abellán and Medina, 2009**; **Abellán et al., 2009**, **2010, 2014; Medina et al., 2019**), including those in which we identified new subdivisions of the central extended amygdala in developing chicken (**Vicario et al., 2014**; see also **Pross et al., 2022**), as well as a new telencephalic subdivision near the frontier with the hypothalamus, related to the subpreoptic region and a ventral part of the medial extended amygdala (**Metwalli et al., 2022**).

## Results

We will start by describing the expression of CRF mRNA in chicken telencephalon throughout development (Figs. 1-5), followed by subsequent descriptions of expression patterns of CRFR1 (Figs. 6-10), CRFR2 (Figs. 11-14) and CRBBP (Figs. 15-18). A summary of the main results is also shown in Tables 3-6.

**Figure 1.**
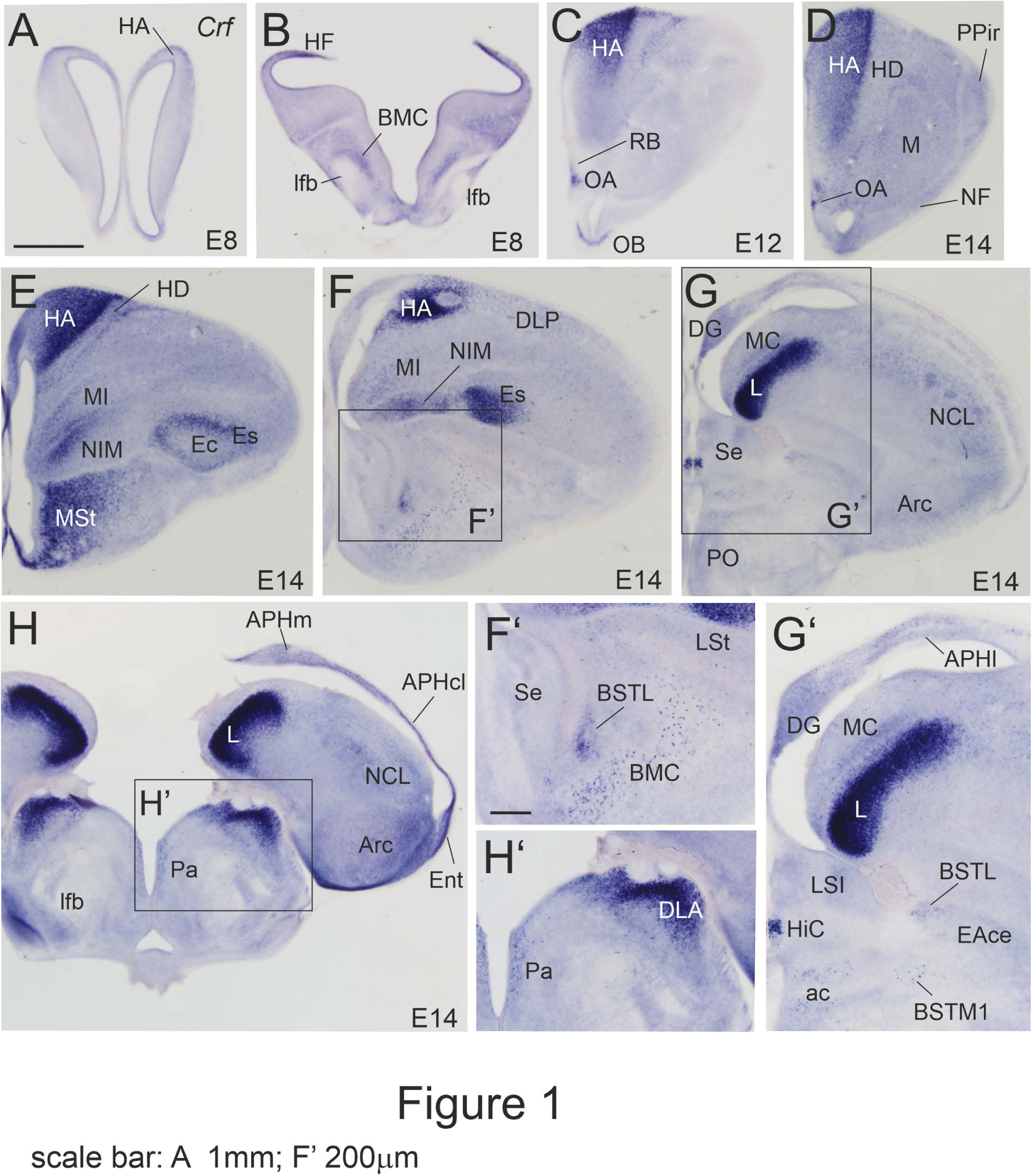
Images of frontal sections through the embryonic telencephalon and anterior hypothalamus of chicken (E8 and E14), from anterior to posterior levels, showing hybridization signal for chicken Crf. F’, G’, and H’ are details of the areas squared in panels F, G and H. Note the strong signal at E14 in visual and auditory pallial areas (hyperpallium, entopallium, field L). Also note the remarkable expression in rostrointermediate parts of the medial striatum, which is in contrast with the low or negligible expression in the lateral striatum and caudal parts of the medial striatum. At this age, expression is also seen in the parvocellular paraventricular hypothalamic nucleus, as well as in several nuclei of the thalamus. See more details in text. For abbreviations see list. Scale bars: A = 1mm (applies to A-H); F’ = 200 μm (applies to F’-H’).

**Table 3:** *Crf* mRNA expression in the chicken telencephalon throughout development

|  | E14 | E18 | P0 | P7 |
| --- | --- | --- | --- | --- |
| DG | +/++ | +/++ | + | + |
| APH | -/+ | +/++ | + | + |
| HA | +++ | ++/+++ | +/+++ | +/+++ |
| IHA | ++++ | ++++ | +++ | ++/+++ |
| HD | + | + | + | + |
| DLP | -/+ | + | + | + |
| MIM - MC | + | + | + | +/++ |
| NIM | +/++ | +/++ | ++ | ++ |
| Es | ++/+++ | +/++ | +/++ |  |
| NCM (Field L) | ++++ | ++++ | ++/+++ | ++/+++ |
| NCIF - NCL | -/+ | + | +/++ | ++ |
| Arc | +/++ | +/++ | +/++ | +/++ |
| OB | + | ++ | ++ | ++/+++ |
| PPir - Pir | -/+ | + | +/++ | ++ |
| MSt - Ac | +/+++ | +/+++ | +/+++ | +/+++ |
| LSt | -/+ | + | + | ++ |
| GP | + | + | + | + |
| VP - SPa | + | + | + | ++ |
| BMC | ++ | +++ | +++ | +++ |
| BSTL | +/++ | +++ | +++ | ++/+++ |
| BSTM | + | + | + | +/++ |
| Ce | -/+ | + | + | +/++ |
| Me | + |  | + | + |
| LS - MS | -/+ | -/+ | -/+ | -/+ |
| HiC - SFi | +/+++ | +/+++ | +/+++ | +/++++ |
| POM | + | + | ++ |  |

**Table 4:** *Crfr1* mRNA expression in the chicken telencephalon throughout development

|  | **E14** | **E18/P0** | **P7** |
| --- | --- | --- | --- |
| **DG** | ++++ | ++++ | +++ |
| **APH** | ++++ | +++ /++++ | ++ /+++ |
| **HA** | + /+++ | ++ /+++ | ++ /+++ |
| **IHA** | ++ /+++ | +++ | +++ |
| **HD** | - /+ | + | ++ |
| **DLP** | ++ | ++ | ++ |
| **M except MVCo** | - /+ | + | ++ |
| **MVCo** | ++ | ++ /+++ | ++ |
| **NF - Bas** | ++ /+++ | +++ | +++ |
| **NI - NIS** | + /++ | +++ | +++ |
| **NCM (Field L)** | ++ | +++ /++++ | +++ |
| **NCIF - NCL** | ++ | +++ | +++ /+++ |
| **Arc** | ++ /++++ | ++ /+++ | ++ /+++ |
| **AO** | ++ | +++ | +++ |
| **PPir - Pir** | ++ /+++ | ++ /++++ | ++ /+++ |
| **Ac** | + /++ | ++ | ++ |
| **MSt - LSt** | - /+ | + /++ | + /++ |
| **GP** | - | + | + |
| **VP - SPa** | - | + | + |
| **BSTL** | - | + | + |
| **BSTM** | - | + | + |
| **Ce** | - | + | + /++ |
| **Me** | + /++ | ++ |  |
| **LS** | + /++ | ++ /+++ | +++ |
| **HiC - SFi** | + /+++ | ++ /+++ | +++ |
| **POM** | ++ | +++ | +++ |
| **SuPO** | ++ | ++ | ++ |

**Table 5:** *Crfr2* mRNA expression in the chicken telencephalon throughout development

|  | **E14** | **E18/P0** | **P7** |
| --- | --- | --- | --- |
| **DG** | ++ | + | + |
| **APH** | + | + | -/+ |
| **HA** | + / ++ | + / ++ | - / + |
| **HD** | ++ / +++ | +++ | ++ |
| **DLP** | + | - / + | - / + |
| **M** | ++ / +++ | ++ / +++ | ++ |
| **Bas** | +++ | + / +++ | + / ++ |
| **NF - NI** | - / + | - / + | - / + |
| **NCM (Field L)** | - | - | - |
| **NCIF - NCL** | + | + | + |
| **Arc** | - / + | - / + | - / + |
| **AO** | + / ++ | + | + |
| **PPir - Pir** | + | + / ++ | + / ++ |
| **MSt** | - | - / + | + |
| **LSt** | + | + | + |
| **GP** | - | - | - |
| **VP - SPa** | - | - / + | - / + |
| **BSTL** | - | - | - / + |
| **BSTM** | - | - / + | - / + |
| **Ce** | - | - / + | - / + |
| **LS - MS** | - / + | - / + | - / + |
| **HiC - SFi** | - | + | + |
| **PO - SuPO** | - / + | - / + | - / + |

**Table 6:** *Crfrbp* mRNA expression in the chicken telencephalon throughout development

|  | **E14** | **E18/P0** | **P7** |
| --- | --- | --- | --- |
| **DG** | + | + | + |
| **APH** | -/++ | -/++ | +/++ |
| **HA** | -/+ | +/++ | +/++ |
| **HD** | - | -/+ | + |
| **DLP** | - | -/+ | + |
| **M** | -/+ | + | + |
| **NF - NI** | -/+ | + | + |
| **Bas – E** | +/++ | -/+ | + |
| **NCM (Field L)** | + | ++ | ++ |
| **NCIF/NCL** | - | + | + |
| **Arc** | -/+ | -/+ | +/++ |
| **PPir/Pir** | - | -/+ | -/+ |
| **MSt** | -/+ | -/+ | + |
| **StC - LSt** | ++/+++ | +/++ | +/++ |
| **GP** | -/+ | -/+ | + |
| **VP/SPa** | -/+ | -/+ | + |
| **pINP** | ++ | + | + |
| **BSTL** | + | -/+ | -/+ |
| **BSTM** | -/+ | -/+ | -/+ |
| **Ce** | -/+ | -/+ | -/++ |
| **LS/MS** | - | -/+ | -/+ |
| **PO - SuPO** | - | - | - |

### Chicken CRF mRNA (c*Crf* or simply *Crf*)

The first embryonic age we studied was E8 (Hamburger and Hamilton stages 34-35 or HH34-35). At this age we already observed moderate *Crf* signal in the primordia of the hippocampal formation (HF, in the medial pallium; mainly its medial part where the dentate gyrus and medial parahippocampal area will form), the apical hyperpallium (HA, in the dorsal pallium), the caudal part of the nidopallium (N, in the ventral pallium) and part of the basal magnocellular complex (BMC, containing corticopetal cells) of the subpallium (Fig. 1A,B).

By E12-E14, as the telencephalon became gradually larger and more mature, the signal intensity was high in some of these areas and *Crf* expression was clearly localized to specific subdivisions (Fig. 1C-G’). In the pallium, the highest expression was seen in the lateral part of the apical hyperpallium (HA), extending to the interstitial nucleus of the hyperpallium (Fig. 1C-E), and in specific parts of the intermediate and caudal nidopallium, including the medial part of the intermediate nidopallium (NIM), the entopallial belt or shell (Es) and the auditory field L (Fig. 1E-G, detail in G’). Light to moderate expression was also observed in the olfactory bulb (including mitral cells) and anterior olfactory area (OB, AO; Fig. 1C,D), parts of the densocellular hyperpallium (HD, Fig. 1E), medial part of the apical hyperpallium (Fig. 1D,E) , several parts of the hippocampal formation (including dentate gyrus [DG], and medial, lateral and caudolateral parahippocampal areas [APHm, APHl, APHcl]; Fig. 1E-G, G’), the entorhinal cortex (Ent; Fig. 1H), parts of the mesopallium (M, mainly its medial intermediate part; Fig. 1E,F), some patches of the caudolateral nidopallium (NCL; Fig. 1G,H), and part of the arcopallium (Arc, including dorsal, core, amygdalopiriform, and amygdalohippocampal areas; Fig. 1H). Very low expression was additionally observed in the prepiriform and piriform cortices (PPir; Fig. 1D), and in nucleus basorostrallis (not shown). In the subpallium of E12-E14 chicken, strong expression of *Crf* was observed in the medial striatum from rostral (including nucleus accumbens) to intermediate levels (MSt; Fig. 1E) and in the large perikarya of the basal magnocellular complex (BMC), which spread over the medial forebrain bundle/ventral pallidum and the medial aspect of the lateral forebrain bundle to reach the intrapeduncular nucleus and medial aspect of the globus pallidus (Fig. 1F, detail in F’). Part of the central extended amygdala also expressed low to moderate levels of *Crf* at E14, mostly in the lateral BST (BSTL; Fig. 1F’,G’). Scattered cells of the medial BST (BSTM) and the medial preoptic region (PO) also expressed moderate levels of *Crf* at this embryonic age (Fig. 1G,G’). In the septum, strong expression was observed in dorsal part of the nucleus of the hippocampal commissure (HiC) and adjacent part of the septofimbrial nucleus, while the medial and latero-intermediate nuclei (LSI) showed light signal (Fig. 1G’).

By E18 (Fig. 2), *Crf* expression pattern in the telencephalon remained quite similar to that seen at E14, with little changes, such as the increase in *Crf* signal in the BSTL and its lateral extension into the perioval zone of the central extended amygdala (Fig. 2C-E, details in C’ and E’), and in the septofimbrial nucleus (SFi; Fig. 2E’). Signal continued to be quite strong in the lateral part of the apical hyperpallium (at E18 became particularly strong in the interstitial nucleus of the apical hyperpallium or IHA; Fig.2A, B, detail in A’), field L of the caudal nidopallium (L; Fig. 2C-E), rostral (accumbens) parts of the medial striatum (MSt; Fig. 2B), and the large perikarya of the basal magnocellular complex (BMC; Fig. 2C, detail in C’). Moderate to strong expression was also observed in part of the olfactory tubercle (Tu, Fig. 2B,C), while light or moderate signal was present in other pallial and subpallial areas. In the pallium, these included the hippocampal formation (including the dentate gyrus [DG] and the parahippocampal areas [APH]; Fig. 2C,D), medial and densocellular parts of the hyperpallium (Fig. 2A), the caudomedial mesopallium (M; Fig. 2A-C), medial part of the intermediate nidopallium (NIM; Fig. 2B), parts of the arcopallium (Arc, Fig. 2E), the prepiriform and piriform cortices (PPir, Pir; Fig. 2A,B), the anterior olfactory area and the olfactory bulb (OB; Fig. 2A). In the subpallium, scattered cells expressing *Crf* were seen at intermediate/caudal levels of the medial and lateral striatum (MSt, LSt), globus pallidus (GP), and intrapeduncular nucleus (INP) (Fig. 2C, detail in C’). The medial aspects of GP and INP also included large perikarya expressing *Crf*, belonging to BMC (Fig. C’). More caudally, scattered *Crf* expressing cells were seen in lateral parts of the central extended amygdala (including peri-intrapeduncular and capsular areas), the diagonal band nucleus and septopallidal area (PaS), part of the septocommissural area (including the caudocentral septal nucleus or CCS), the BSTM and the preoptic region (Fig.2D,E,E’).

**Figure 2.**
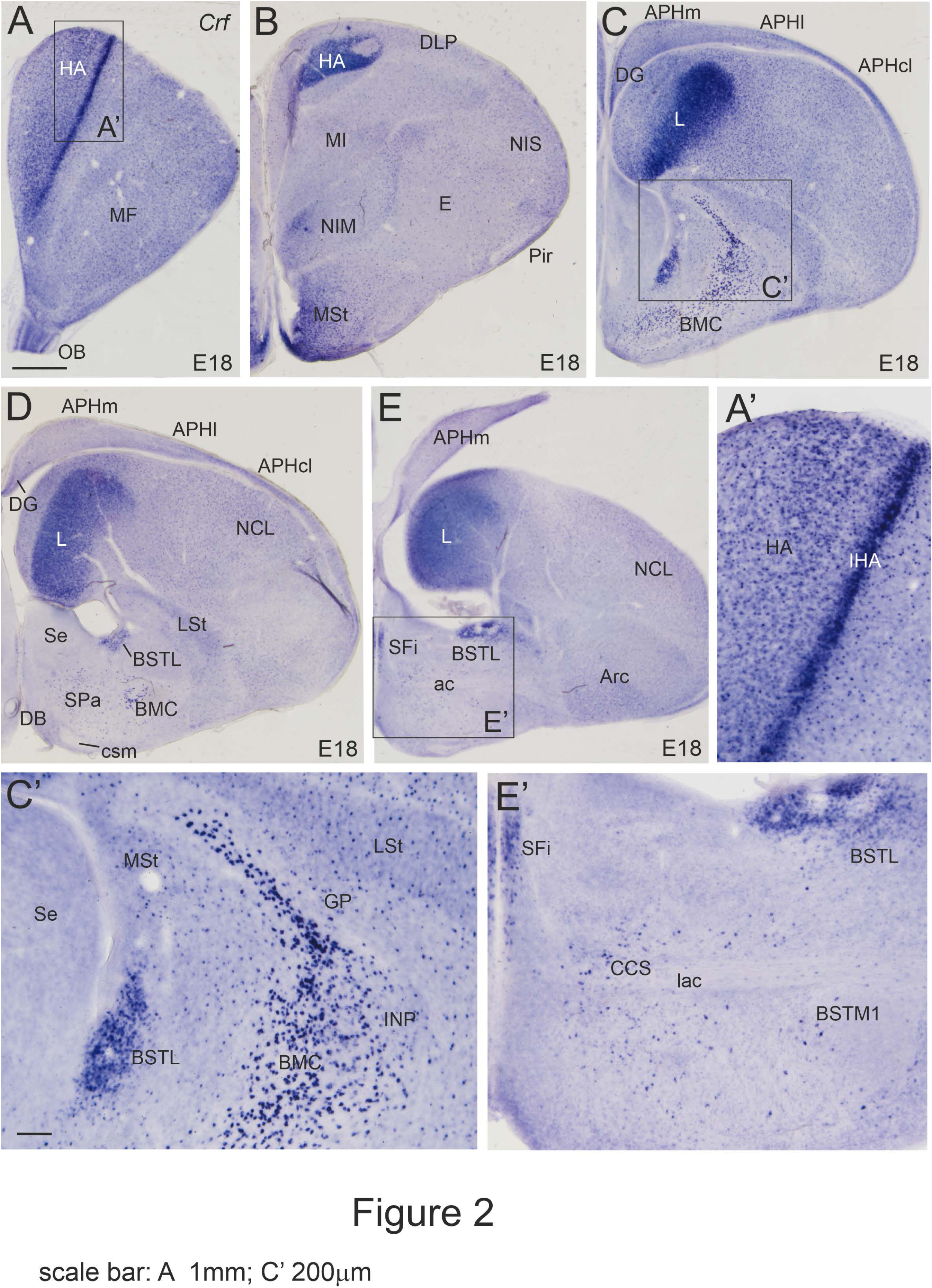
Images of frontal sections through the embryonic telencephalon of chicken (E18), from anterior to posterior levels, showing hybridization signal for chicken Crf (details in A’, C’ and E’, from the squared areas in panels A, C, and E). For abbreviations see list. Scale bars: A = 1mm (applies to A-E); C’ = 200 μm (applies to A’, C’, E’).

At hatching (P0), the *Crf* expression pattern remained quite similar to that seen at E18, although the intensity of the expression became more homogeneous between areas, with a tendency to be light to moderate. The only sites where the signal was moderate to high were the IHA in the hyperpallium (Fig. 3A), field L in the caudal nidopallium (Fig. 3C,D), part of the piriform cortex (Pir; Fig. 3C), the septofimbrial nucleus (SFi, just dorsal to HiC; Fig. 3C, detail in C’), the rostromedial striatum (MSt, including nucleus accumbens; Fig. 3A), the BSTL (Fig. 3C, detail in C’), the perikarya of the basal magnocellular complex (BMC; Fig. 3B), and the medial preoptic region (PO; Fig. 3C). At P7, the pattern remained similar, and most of the same sites continued to show higher levels of *Crf* expression than the surrounding areas (Figs. 4B,C,E; 5B-D). In addition, the olfactory bulb and the anterior olfactory area also showed moderate to high expression at P7 (Fig. 4A). In the olfactory bulb, many cells of the mitral cell layer and some of the granular cell layer expressed high levels of *Crf*. Except the olfactory bulb, olfactorecipient areas, hyperpallium, medial intermediate nidopallium and field L, in the rest of the pallium, *Crf* expressing cells were less densely grouped at P7 compared to previous ages (Figs. 4, 5). In the hippocampal formation, only few, scattered cells were observed in the medial layer of the V-field (containing the dentate gyrus; DG) and the medial and lateral parts of the parahippocampal area (APHm, APHl; Fig. 5A). In the subpallium, the distribution and density of *Crf* expressing cells was quite similar between P7 and P0. Expression continued to be quite high in many cells of SFi and adjacent dorsal aspect of HiC, BSTL and BMC (Fig. 5B-D). Some cells expressing *Crf* were also present in the septopallidal area, lateral striatum, globus pallidus and lateral parts of the central extended amygdala, where a distinct group of *Crf* expressing cells became visible in the medial aspect of the capsular central amygdala (Fig. 5C).

**Figure 3.**
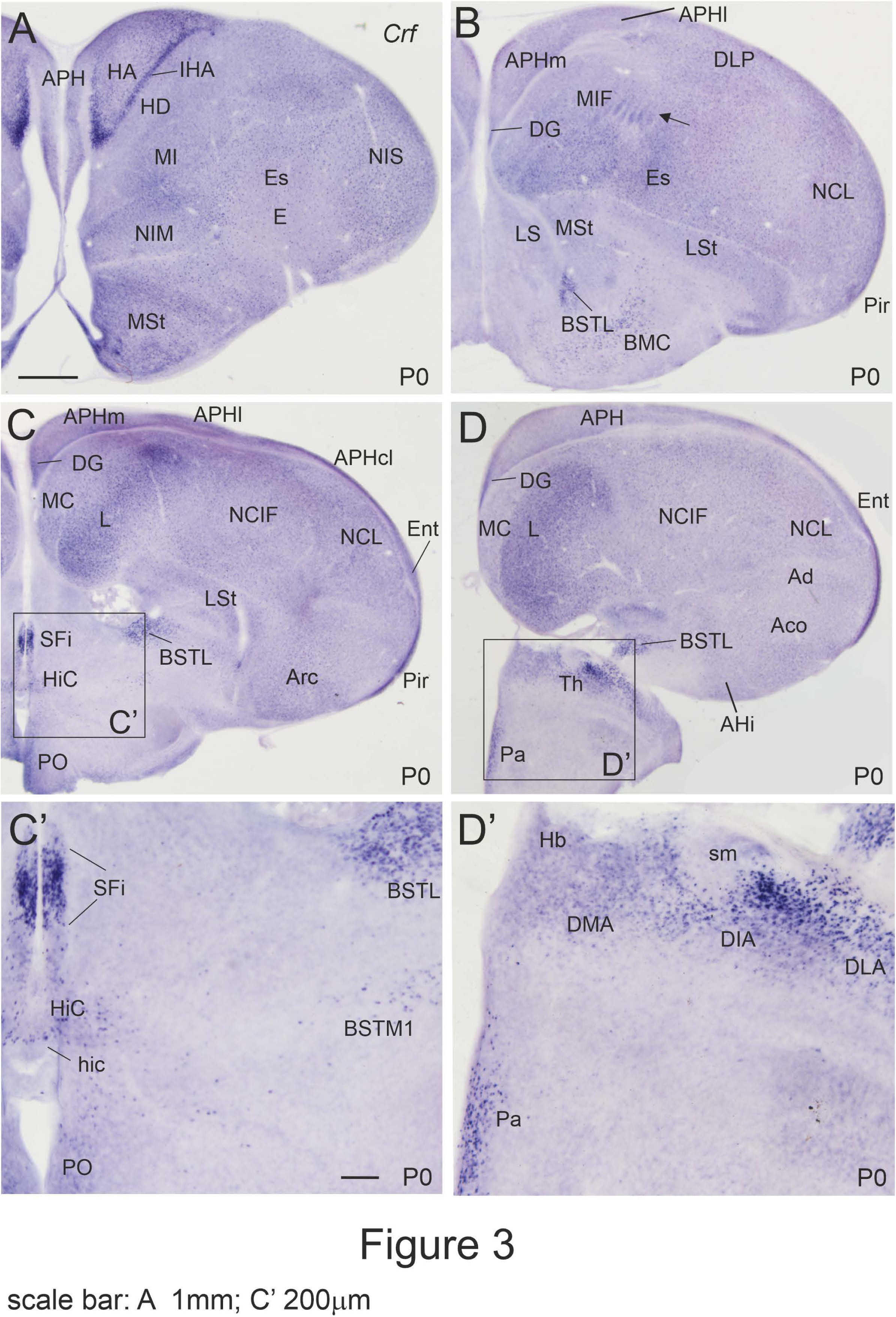
Images of frontal sections through the telencephalon (A-D, C’) and anterior hypothalamus (D, D’) of chicken at hatching (P0), from anterior to posterior levels, showing hybridization signal for chicken Crf (details in C’ and D’, from the squared areas in panels C and D). For abbreviations see list. Scale bars: A = 1mm (applies to A-D); C’ = 200 μm (applies to C’, D’).

**Figure 4.**
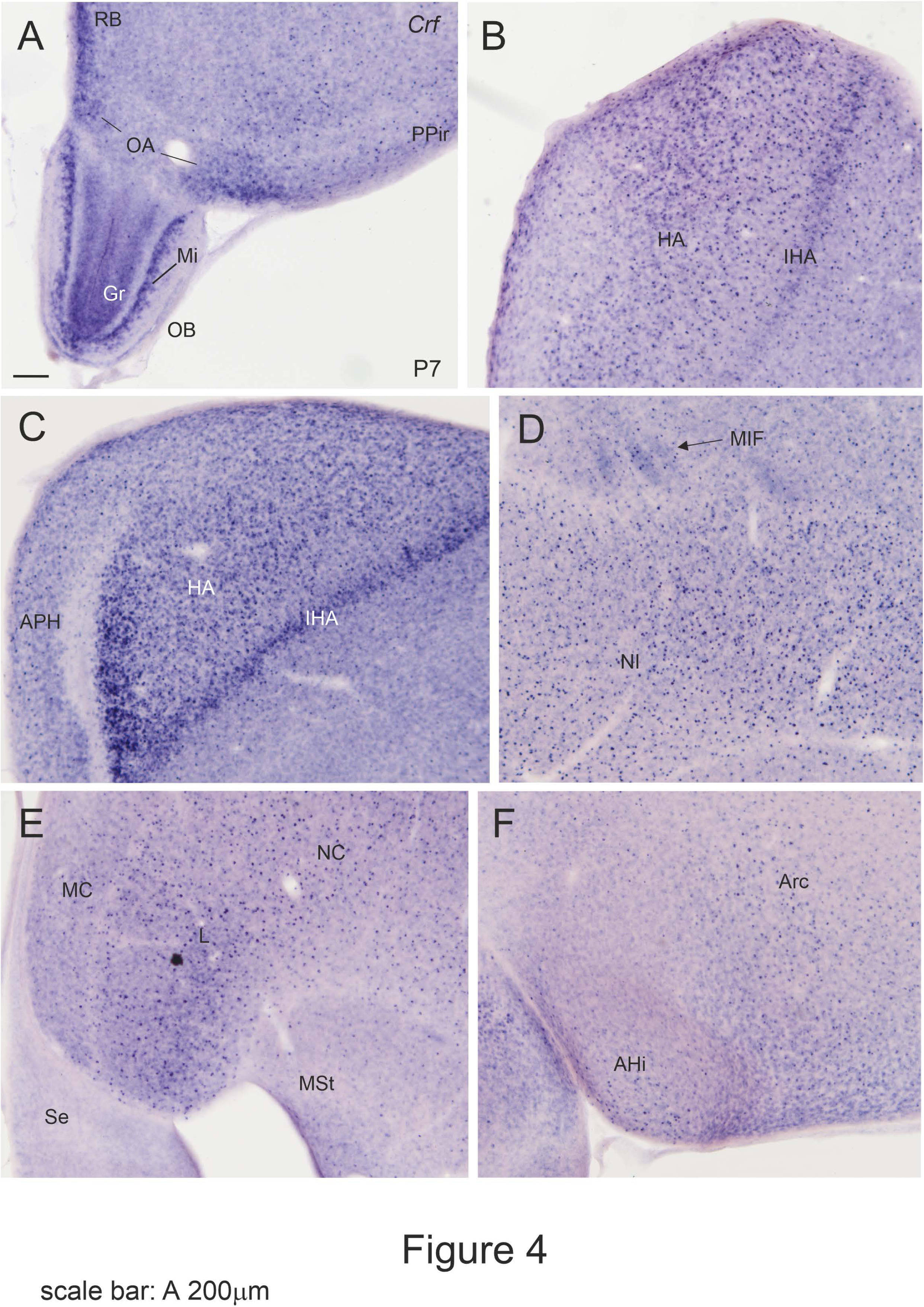
Details of frontal sections through the telencephalon of chicken at P7, showing hybridization signal for chicken Crf. A is at the level of the olfactory bulb and anterior olfactory area; B and C show details of the hyperpallium, at different anteroposterior levels (C is more posterior and also shows part of the parahippocampal formation); D is a detail of the intermediate nidopallium, and the patches of the mesopallial island field near the mesopallium-nidopallium boundary (arrows); E shows the caudomedial nidopallium and mesopallium; F is a detail of the arcopallium. For abbreviations see list. Scale bars: A = 200 μm (applies to all).

**Figure 5.**
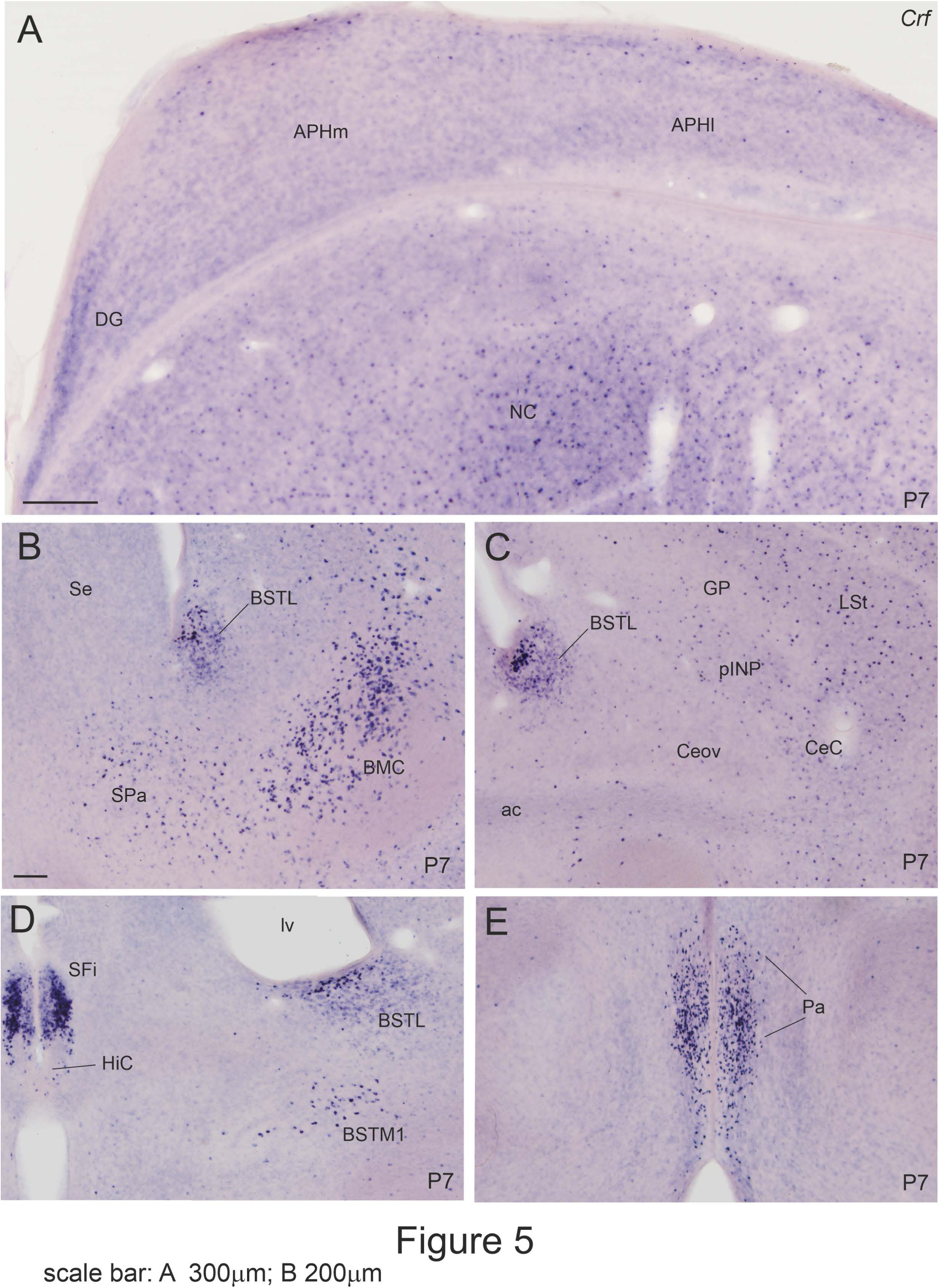
Details of frontal sections through the telencephalon (A-D) and anterior hypothalamus (E) of chicken at P7, showing hybridization signal for chicken Crf. A is at the level of the hippocampal formation; B-D show details of the subpallium, from anterior to posterior levels, showing the basal magnocellular complex (BMC), several nuclei of the extended amygdala (incuding BSTL, BSTM, pINP and CeC); D also shows the septum, with strong signal in HiC/SFi nuclei. E shows a detail of the parvocellular part of the paraventricular hypothalamic nucleus. For abbreviations see list. Scale bars: A = 200 μm (applies to all).

Outside the telencephalon, we would like to remark the presence of cells expressing moderate levels of *Crf* in the parvocellular part of the paraventricular hypothalamic nucleus, on the top of the hypothalamo-pituitary-adrenal axis, from E14 (Fig. 1H, detail in H’). Compared to E14, the expression levels in these cells appeared to be higher at E18 and hatching (P0; Fig. 3D, detail in D’), and even higher at P7 (Fig. 5E), at least based on hybridization signal. Many *Crf* expressing cells were also seen in visual, auditory and somatomotor parts of the thalamus from E14 (Fig. 1H, detail in H’). In particular, strong expression was seen in many cells of the anterior dorsolateral/dorsointermediate nuclei (DLA, DIA). In addition, some *Crf* expressing cells were present in the dorsointermediate ventral anterior nucleus,), ventrointermediate nucleus, suprarotundus (or epirotundus) and subrotundus nuclei, medial pole of nucleus rotundus, and periovoidal region or shell). In the dorsal midbrain, visual and auditory centers (optic tectum and torus semicircularis) also showed *Crf* expression at least from E14.

### Chicken CRF receptor 1 mRNA (*cCrfr1 or Crfr1*)

At E8, expression of *Crfr1* was remarkably strong or very strong in parts of the pallium, including the primordium of the hippocampal formation, several olfactorecipent areas at the surface of the ventral pallium (including the primordium of the piriform cortex), the intermediate and caudal levels of the nidopallium, and the arcopallium (Fig. 6A-C). At this age, moderate expression was also present in the dorsolateral pallium. In contrast, the subpallium was quite poor in *Crfr1* expression, except for two small groups of subpial cells at the septofimbrial nucleus (medially; Fig. 6A) and the olfactory tubercle (laterally, arrow in Fig. 6A), and few scattered cells in the lateral striatum, the nucleus of the lateral olfactory tract (LOT) and the preoptic region (PO) (Fig. 6B).

At E14, the telencephalon was more developed and it was possible to identify better the areas expressing the receptor mRNA (Fig. 6D-I). *Crfr1* expression continued to be moderate to high in most subdivisions of the pallium, except the mesopallium, but a clear pattern emerged with areas of high expression and others with light or no expression. Both the hippocampal complex (including the hippocampal formation and the entorhinal cortex) and the arcopallium showed the highest and most extensive expression levels (Fig. 6G-I). In the hippocampal complex, high expression covered all areas (DG, APH, Ent) and layers, along all mediolateral and anteroposterior levels (Fig. 6E-I). In the arcopallium, moderate to strong expression was seen in many areas, increasing from anterior to posterior levels (Fig. 6G-I), and the strongest signal was observed in the dorsal, amygdalopiriform, and amygdalohippocampal area (Fig. 6I). All the dorsolateral pallium (DLP) (Fig. 6F) and all olfactorecipient areas of the ventral pallium (anterior olfactory area [OA; Fig. 6D], prepiriform cortex [PPir; Fig. 6D, F] and piriform cortex [Pir; Fig. 6G,H]) showed moderate to high expression. In the hyperpallium and nidopallium, expression was high in some areas and low in others, as follows. In the hyperpallium, moderate to high expression was observed medially in the apical hyperpallium and in the IHA, but expression was low in the lateral apical and densocellular hyperpallium (Fig. 6D,E). In the nidopallium (N), signal was moderate to strong in the frontal, part of the intermediate, and part of the caudal nidopallium (Fig. 6D-I). At frontal levels (NF), expression was moderate to high in the frontolateral nidopallium (NFL) and in nucleus basorostralis (Bas) (Fig. 6D,E). At intermediate levels (NI), expression was moderate in the superficial stratum (NIS), but the entopallium and the medial area only contained light or negligible expression (Fig. 6E,F). In the caudal nidopallium (NC), moderate expression was observed in the caudolateral area (NCL), the dorsal nidopallium, and the periphery of field L in the caudomedial area (Fig. 6G,H).

**Figure 6.**
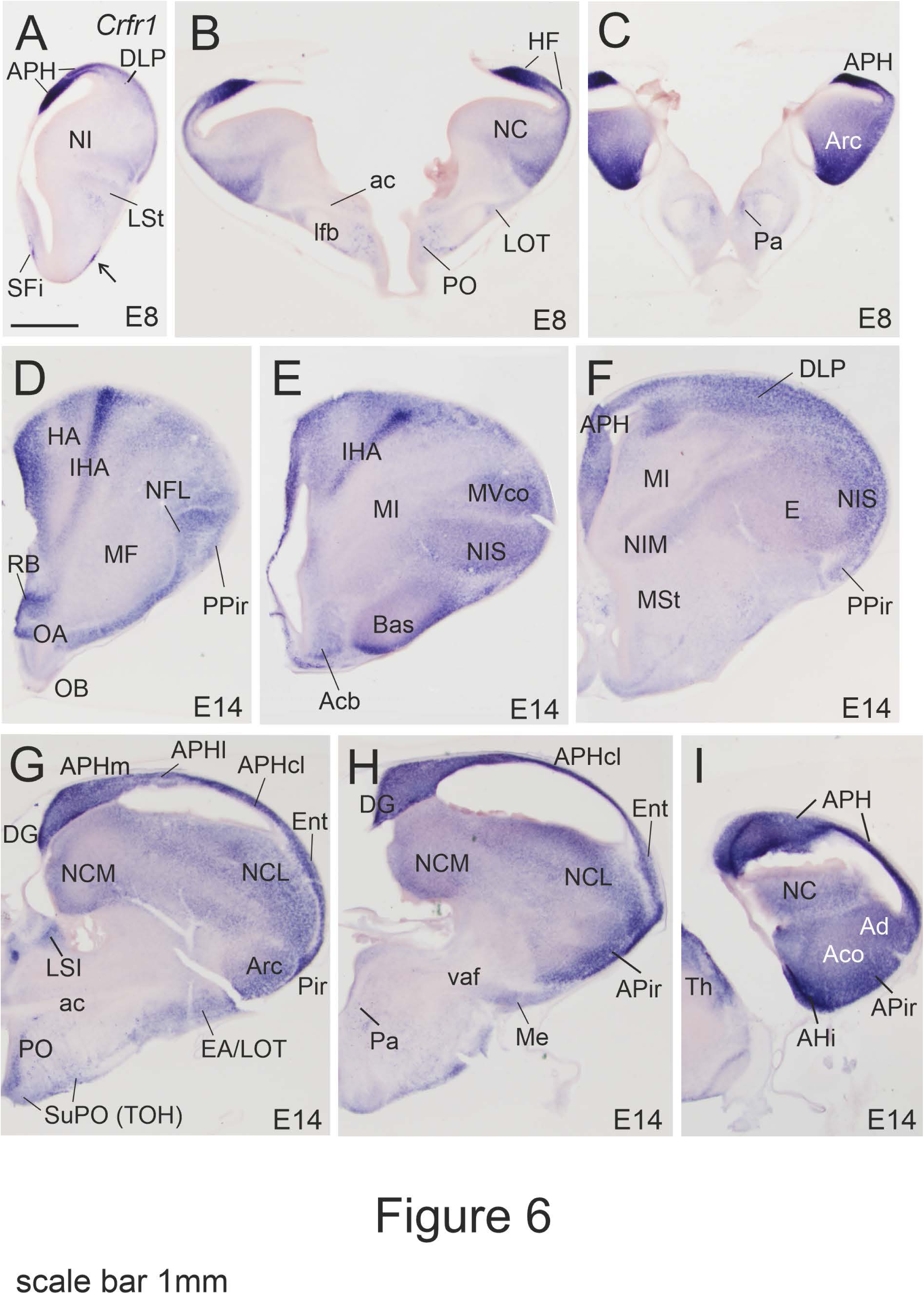
Images of frontal sections through the embryonic telencephalon and anterior hypothalamus of chicken (E8 and E14), from anterior to posterior levels, showing hybridization signal for chicken Crfr1. Note the strong signal from E8 in the hippocampal formation and in pallial areas involved in high-order association (MVCo, NCL) and sensorimotor integration (HA, Arc). From E8, expression is also seen in the septum (including SFi), in a subpial patch of the olfactory tubercle (arrow in panel A), and in the paraventricular hypothalamic nucleus. See more details in text. For abbreviations see list. Scale bars: A = 1mm (applies to all).

Regarding the mesopallium, this pallial subdivision was quite poor in *Crfr1* expression, except for its frontomedial, retrobulbar pole (RB, Fig. 6D), and the core nucleus of the ventral mesopallium (MVco, Fig. 6E). In contrast to the pallium, at E14 the subpallium was almost free of *Crfr1* expression, except part of nucleus accumbens (Fig. 6E), part of the septum (laterointermediate septum [LSI] and the part of the septofimbrial nucleus [SFi] adjacent to HiC; Fig. 6G), the medial amygdala (Me; Fig. 6H), and the medial preoptic nucleus (Fig. 6G), which showed moderate levels of expression. Ventral to the subpallium, moderate expression was also present in the subpreoptic region (SuPO) of the telencephalon-opto-hypothalamic domain or TOH (Fig. 6G). The cell corridor extending between the arcopallium and the TOH, which includes the nucleus of the lateral olfactory tract (LOT), also contained moderate expression of *Crfr1* at E14 (Fig. 6G).

At E18, *Crfr1* expression intensity increased in the pallium, and it spread to cover more areas. At this age, all areas of the hippocampal complex, apical hyperpallium, nidopallium, and arcopallium showed moderate to high levels of *Crfr1* (Fig. 7A-F). In the hyperpallium, the highest expression was seen in the apical hyperpallium and IHA (Fig. 7A), while in the nidopallium, the highest signal was observed in the frontolateral (NFL; Fig. 7A) and medial intermediate (NIM; Fig. 7B,C) areas, as well as in field L, the patches of the island field of the caudal nidopallium (NCIF), and the caudolateral nidopallium (NCL) (Fig. 7E,F). In comparison, most of the densocellular hyperpallium (HD, except its superficial part) and most parts of the mesopallium (M, except MVco) showed light expression (Fig. 7A-C). Close to the pial surface of the ventral pallium, the piriform cortex also showed strong expression of *Crfr1* (Fig. 7D,E). In the subpallium, expression remained relatively low, although the signal slightly increased in some areas, such as the striatal capsule (StC; Figs. 7D, 8A), the accumbens shell (AcS; Fig. 7B,C), the globus pallidus (GP; Fig. 8A), the intrapeduncular nucleus (INP; Fig. 8A), part of the central extended amygdala, including the oval nucleus and peri/post-intrapeduncular island field (Ceov, pINP; Figs. 7E,F; 8B), the septum (including the lateral septum [LS] and the septofimbrial nucleus) (Figs. 7E; 8A), and the medial preoptic region (PO; Fig. 7F, 8C). At E18, scattered *Crfr1* expressing cells were also seen in the medial BST of the medial extended amygdala (Fig. 7F, detail in F’), and in other striatal and ventral pallidal parts of the basal ganglia not mentioned before (Fig. 8A).

**Figure 7.**
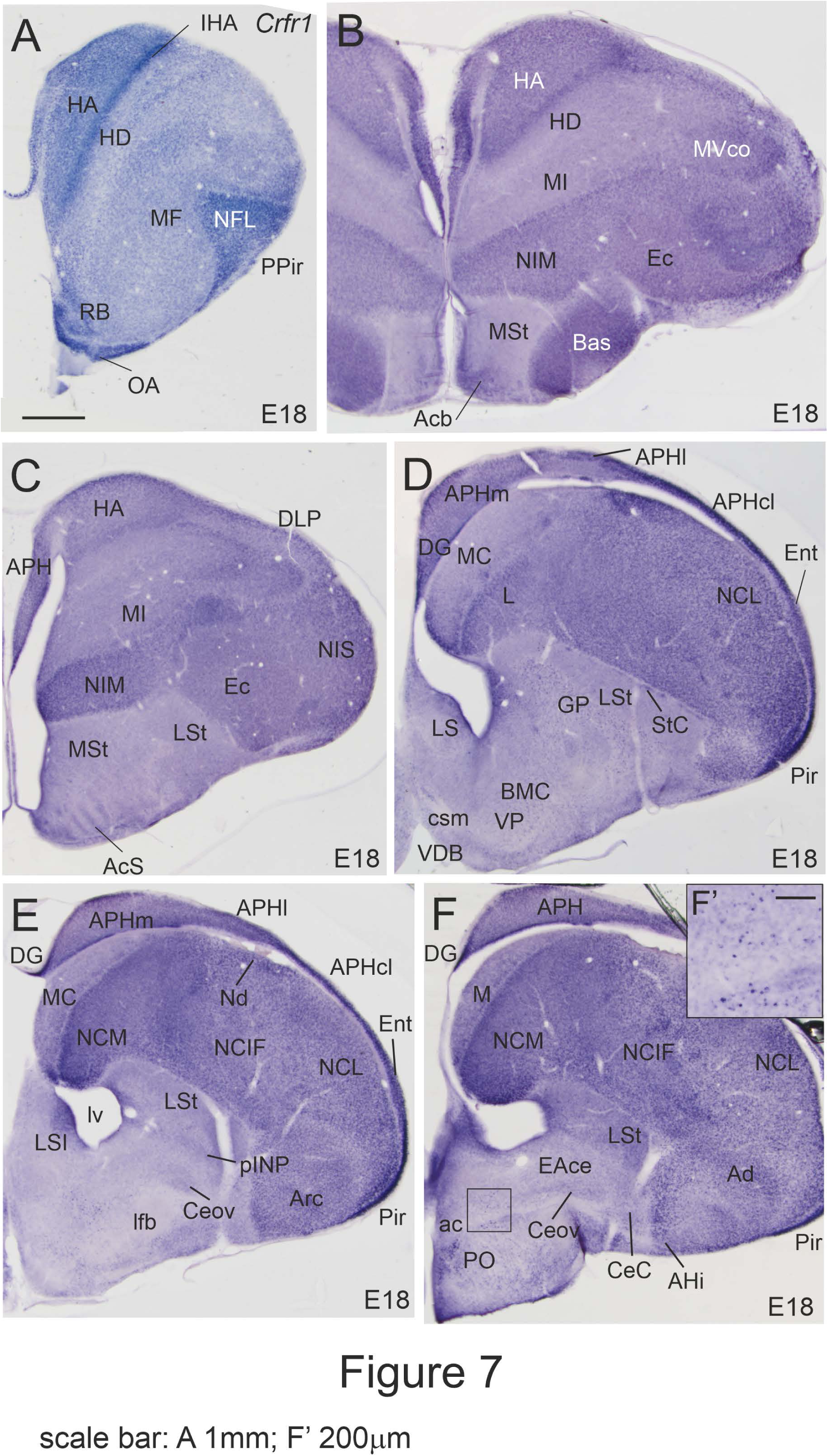
Images of frontal sections through the embryonic telencephalon and anterior hypothalamus of chicken (E18), from anterior to posterior levels, showing hybridization signal for chicken Crfr1. Note the strong signal in the pallium, but low expression in the subpallium, except parts of the striatum, septum, and medial preoptic region. The detail in F’ shows scattered cells in BSTM (from the squared areas in panel F). See more details in text. For abbreviations see list. Scale bars: A = 1mm (applies to A-F); F’ = 200 μm.

**Figure 8.**
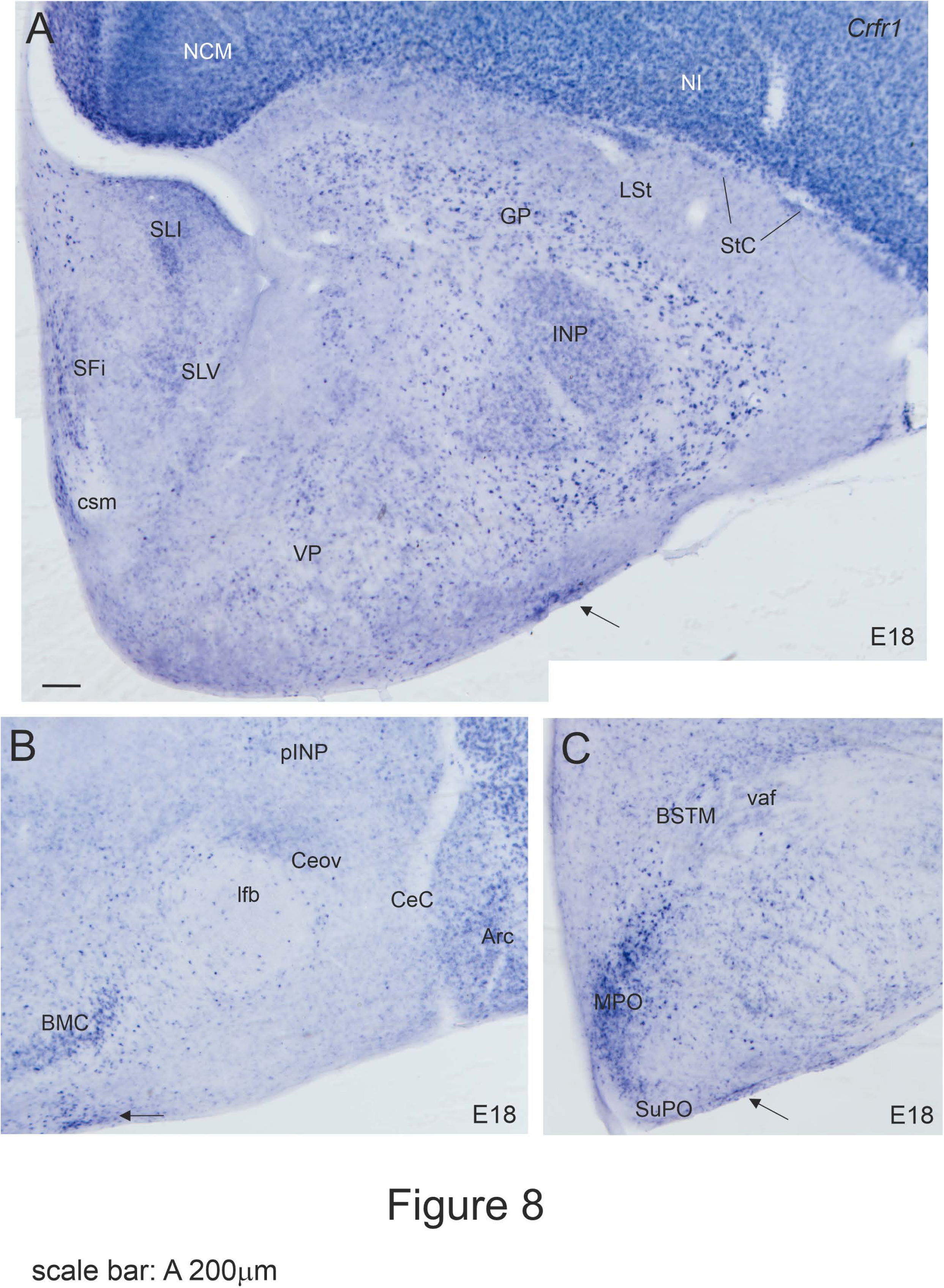
Details of Crfr1 in the subpallium of the embryonic telencephalon at E18. A is a photomontage. Compared to the pallium, expression in the subpallium is generally low, with a few exceptions. Note the signal in the striatal capsule (StC), globus pallidus, INP, parts of the septum, and medial preoptic region. The arrows in A-C) point to subpial expression that appears to extend from the subpreoptic region (SuPO, in the TOH domain; C), reaching the lateral subpallial surface, where the olfactory tubercle locates (A). Low signal is also seen in parts of the extended amygdala (Ceov, BSTM). See more details in text. For abbreviations see list. Scale bars: A = 200 μm (applies to all).

The expression pattern of *Crfr1* remained similar at hatching. One week later (P7), the pattern remained the same, but the signal intensity increased in parts of both the pallium and the subpallium (Figs. 9, 10). In the pallium, expression increased in the mesopallium, while keeping a lower intensity compared to other pallial subdivisions, such as the apical hyperpallium and the nidopallium (Fig. 9B,C). In the subpallium, expression was a bit stronger in the septum (Fig. 9D), parts of the extended amygdala (Figs. 9C,C’; 10), and the preoptic region (Figs. 9D; 10). In the central extended amygdala, moderate expression was visible in the oval nucleus (Ceov), part of the peri/post-intrapeduncular island field (pINP) (Figs. 9C, detail in C’; 9D, detail in 10), and the capsular central amygdala (Fig. 10). In the septum, moderate to high signal was seen in the laterointermediate nucleus (LSI), septofimbrial nucleus (SFi), and nucleus of the hippocampal commissure (HiC) (Figs. 9D, detail in 10). Scattered cells expressing *Crfr1* were also observed in nuclei of the septopreoptic territory, including the septocommissural nucleus or CoS (Fig. 10). In the preoptic region, high expression was observed in the medial preoptic nucleus, although scattered cells expressing *Crfr1* were also seen in the medial preoptic area (Fig. 10). Finally, the subpreoptic region (SuPO, in TOH) and the subpial cell corridor extending from TOH to the amygdala contained abundant cells expressing *Crfr1* (arrows in Fig. 10). Several cell populations with *Crfr1* expression were observed along this subpial corridor, including a migrated group tentatively identified asthe supraoptic nucleus (SO) (Fig. 10).

**Figure 9.**
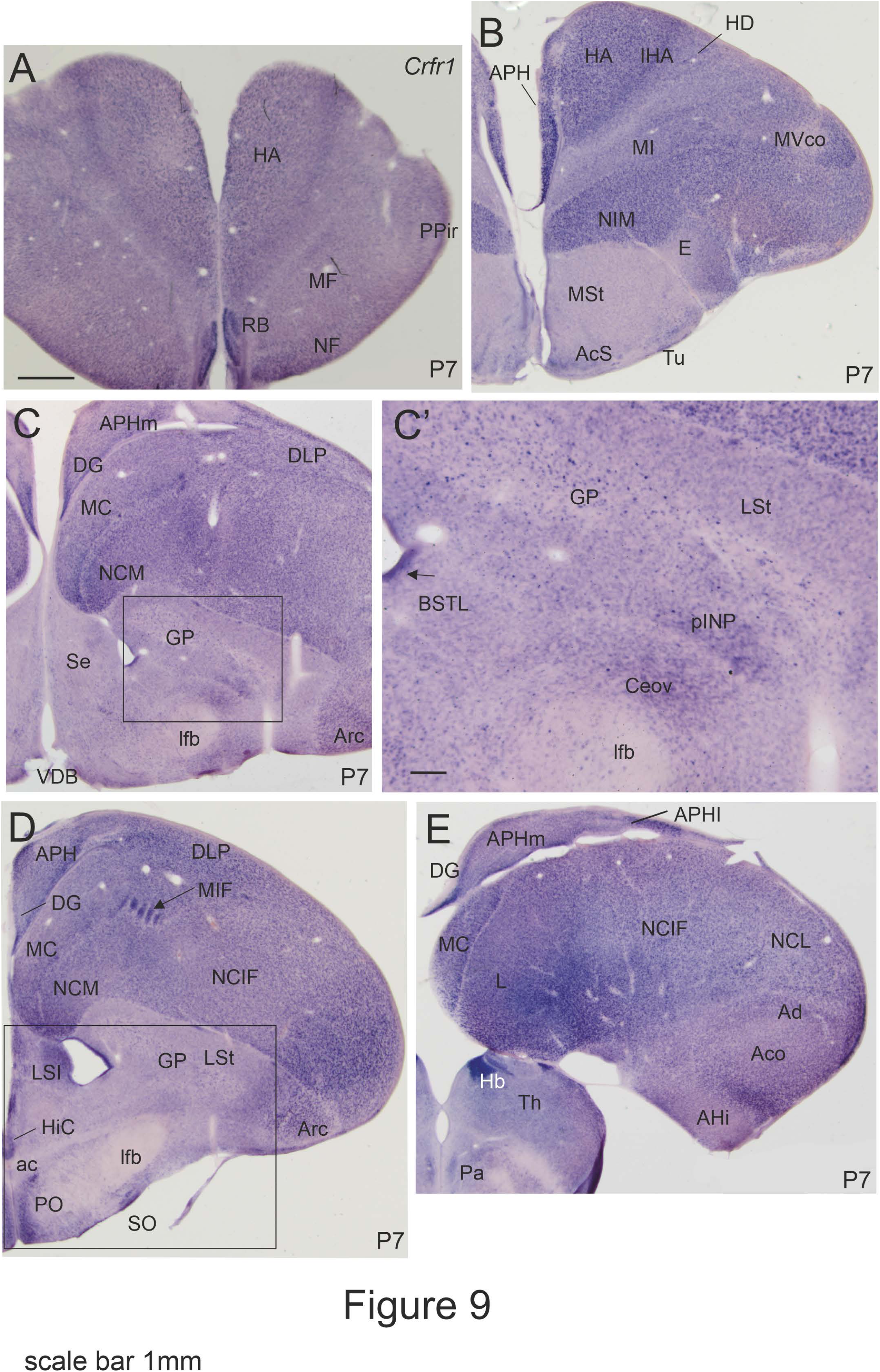
Images of frontal sections through the telencephalon and anterior hypothalamus of chicken at P7, from anterior to posterior levels, showing hybridization signal for chicken Crfr1. C’ shows a detail of the area squared in C. The squared area in D is shown in Figure 10 at higher magnification. Note the strong signal in the pallium, but low to moderate expression in the subpallium. In the subpallium, moderate signal is observed in part of the striatum, septum, central extended amygdala (Ceov, pINP) and medial preoptic region. See more details in text. For abbreviations see list. Scale bars: A = 1mm (applies to A-C, D, E); C’ = 200 μm.

**Figure 10.**
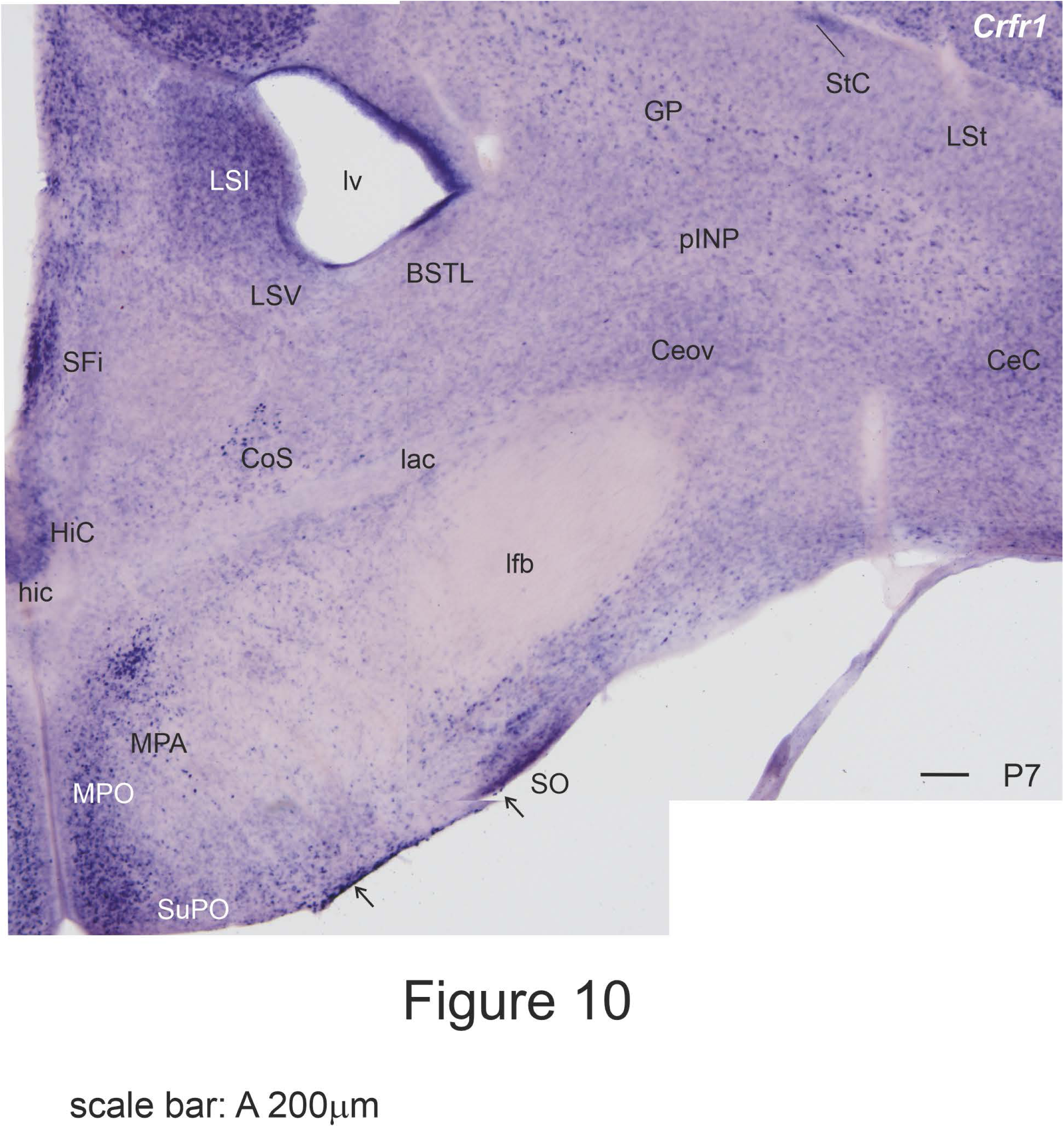
Detail of Crfr1 in the subpallium of chicken at P7 (photomontage). Note the signal in the striatal capsule (StC), globus pallidus, parts of the extended amygdala (pINP, Ceov, CeC) and septum (LSI, HiC, SFi), and medial preoptic region. Expression is also seen in a subpial area (aroows) that appears to extend dorsolaterally from the subpreoptic region (SuPO, in the TOH domain), reaching the lateral surface of the subpallium. Part of the supraoptic nucleus (SO) appears to locate in the middle of this Crfr1 expressing subpial area. Low signal is also seen in parts of the extended amygdala (Ceov, BSTM). See more details in text. For abbreviations see list. Scale bars: A = 200 μm.

### Chicken CRF receptor 2 mRNA (*cCrfr2 or Crfr2*)

Throughout development, *Crfr2* showed an expression pattern remarkably different from that of *Crfr1*. At E8, *Crfr2* signal was very high in the developing mantle of the mesopallium (lateral pallium), in medial and intermediate parts of the hippocampal formation, and in nucleus basorostralis of the nidopallium (Fig. 11A). Light to moderate expression was also seen in the lateralmost part of the developing hippocampal formation, the mantle of the dorsal hyperpallium and dorsolateral pallium, the medial part of the intermediate nidopallium, and in a superficial layer of the caudal ventral/ventrocaudal pallium (Fig. 11A,B). The rest of the pallium was basically free of expression. At E8, the subpallium was free of *Crfr2* expression, while signal was visible in the medial part of the subpreoptic region, in TOH (Fig. 11B).

**Figure 11.**
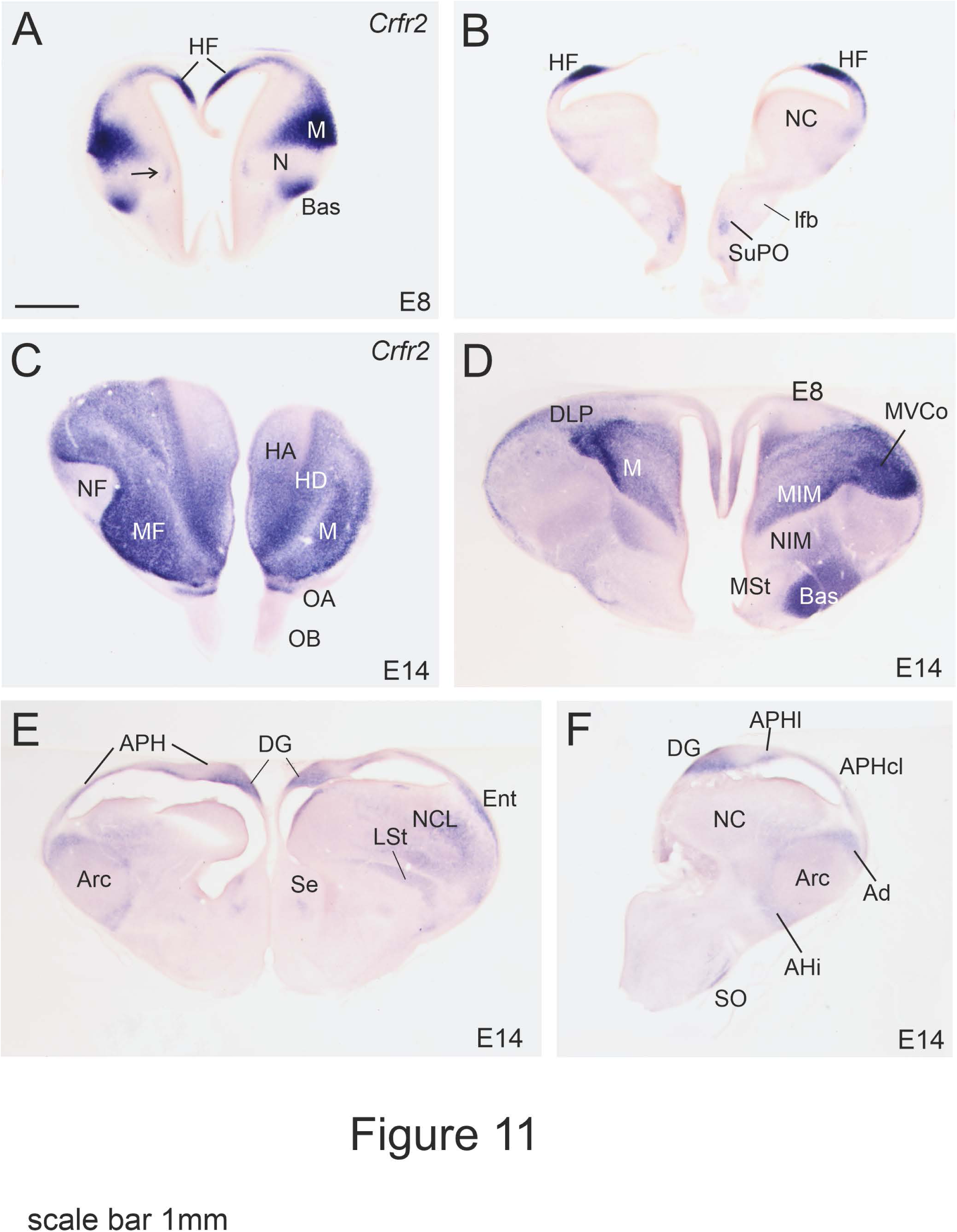
Images of frontal sections through the embryonic telencephalon and anterior hypothalamus of chicken (E8 and E14), from anterior to posterior levels, showing hybridization signal for chicken Crfr2. Note the strong signal from E8 in the hippocampal formation, the mesopallium (M), and in nucleus basorostralis (Bas). Expression in the hippocampal formation becomes less intense at E14. See more details in text. For abbreviations see list. Scale bars: A = 1mm (applies to all).

At E14, *Crfr2* expression remained high in most parts of the mesopallium, being particularly strong in MVco (Fig. 11C,D). At this age, *Crfr2* was also high in the densocellular hyperpallium (HD; Fig. 11C), while the dorsolateral pallium and apical hyperpallium showed moderate expression only in part of the area (Fig. 11C,D). The nidopallium and arcopallium showed only light or no expression (Fig. 11C-F), the only exception being the nucleus basorostralis, which maintained high expression at E14 (Fig. 11D). In the rest of the nidopallium, light expression was visible in part of the medial intermediate nidopallium (just medial to the entopallium) (Fig. 11D), and part of the caudolateral nidopallium (Fig. 11E). In the arcopallium, light expression was present in the dorsal arcopallium and the amygdalohippocampal area (Fig. 11F). In the hippocampal formation, expression was light or moderate in several areas from rostromedial to caudolateral, including the primordia of the dentate gyrus and part of the medial, lateral and caudolateral parahippocampal areas (Fig. 11D-F). The entorhinal cortex (Fig. 11E) and the anterior olfactory area (Fig. 11C) also displayed moderate expression. Regarding the subpallium, very light expression was seen in the lateral striatum (Fig. 11E), and lateral and ventrointermediate parts of the septum (Fig. 11E). More ventrally, in the subpreoptic region, signal was seen in a superficial area that appear to correspond to the supraoptic nucleus (Fig. 11F).

At E18, the expression pattern of *cCrfr2* remained quite similar to that seen at E14 (Fig. 12). Expression was generally high in the mesopallium, but now it was possible to recognize differences in signal intensity between subdivisions within this pallial division (Fig. 12A-E). The highest intensity was seen in the ventral mesopallium, especially in its known associative centers of the ventral mesopallium at frontal (MF; Fig. 12A), intermediate (MVCo-also called MVL-and MVM; Fig. 12B,C), and caudal levels (MC, Fig. 12D,E), including the mesopallial island field (MIF) adjacent to the boundary with the nidopallium (Fig. 12A,C,D). The dorsal mesopallium (MD) contained areas of high or moderate expression and areas of light expression (Fig. 12B,C). In the hyperpallium, expression was high in the densocellular subdivision (HD), but signal in the apical hyperpallium (HA) ranged from moderate at rostral levels (Fig. 12A) to low or negligible at intermediate or caudal levels (Fig. 12B,C). At this age, *Crfr2* expression in the hippocampal formation, dorsolateral pallium, nidopallium and arcopallium was light (for example, in DG and NCL) or negligible in most areas. Light expression was also observed in the olfactory bulb, anterior olfactory area (Fig. 12A), prepiriform cortex (Fig. 12A), and moderate to high expression was seen in the piriform cortex (Fig. 12C-E), and the entorhinal cortex (Fig. 12E). In the subpallium, signal was low to moderate in the lateral striatum, lateral part of the medial striatum, part of the olfactory tubercle, and lateral part of the peri/post-intrapeduncular island field (Fig. 12D), and extremely light in the BSTM and the medial part of the preoptic region (Fig. 12D,E). In the septum, light expression was visible in the lateral and ventrointermediate subnuclei (Fig. 12C), and in the nucleus of the hippocampal commissure (Fig. 12D,E).

**Figure 12.**
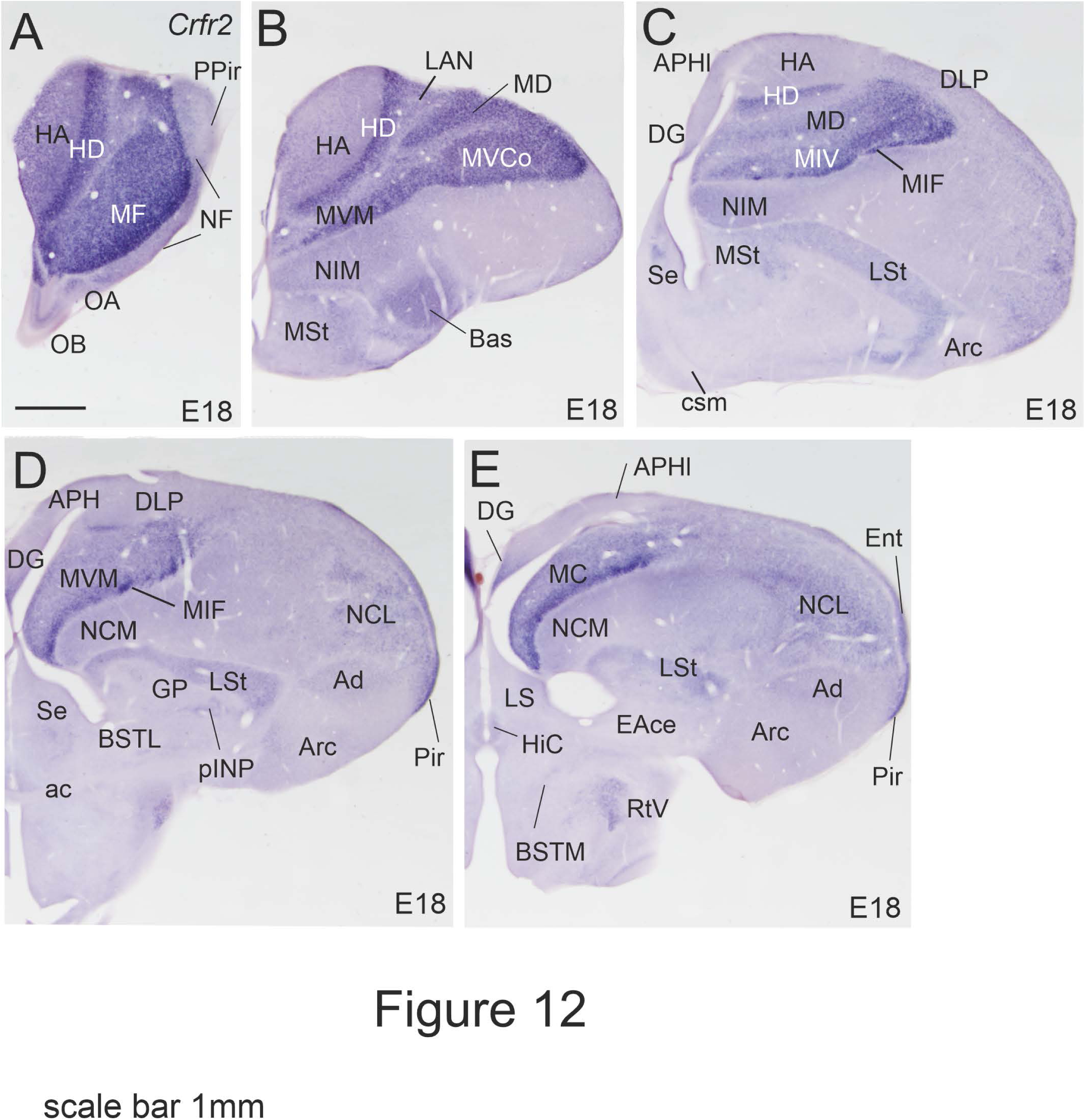
Images of frontal sections through the embryonic telencephalon and anterior hypothalamus of chicken (E18), from anterior to posterior levels, showing hybridization signal for chicken Crfr2. Note the strong signal in the mesopallium (M) and densocellular hypepallium (HD). The olfactorecipient piriform cortex (Pir) also shows expression. See more details in text. For abbreviations see list. Scale bars: A = 1mm (applies to all).

At P0 (Fig. 13), the pattern remained similar to that at E18, but signal intensity increased in the dentate gyrus of the hippocampal formation, to reach levels comparable to those in the mesopallium (Fig. 13D-F, detail in F’’). At this age, expression was also intense in the piriform cortex (Fig. 13F, detail in F’). In the subpallium, in addition to other areas mentioned above for E18, light signal was now visible in other parts of the medial striatum, including patches related to the shell of nucleus accumbens (AcS; Fig. 13C). In addition, light to moderate signal was seen not only in the lateral striatum (LSt), but also in the adjacent striatal capsule (StC, Fig. 13C,D). By P7, the intensity of *Crfr2* expression generally decreased, while the pattern was similar to that seen at hatching and near hatching ages (Fig. 14).

**Figure 13.**
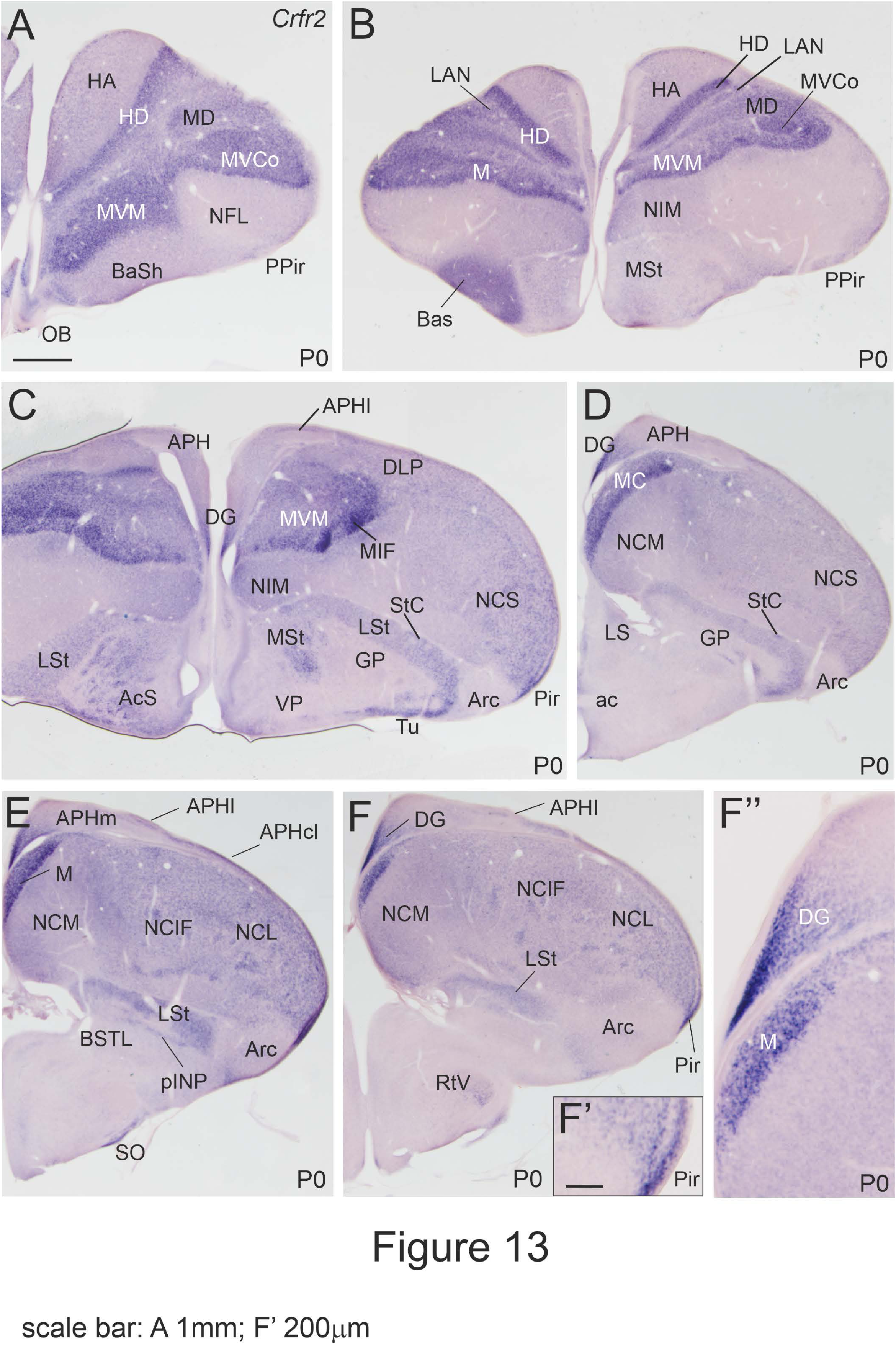
Images of frontal sections through the telencephalon and anterior hypothalamus of chicken at hatching (P0), from anterior to posterior levels, showing hybridization signal for chicken Crfr2. Note the strong signal in the mesopallium (M) and densocellular hypepallium (HD). F’ shows a detail of the intense signal in the piriform cortex (Pir), while F’’ shows a detail of the signal in the dentate gyrus (DG) of the hippocampal formation. See more details in text. For abbreviations see list. Scale bars: A = 1mm (applies to all).

**Figure 14.**
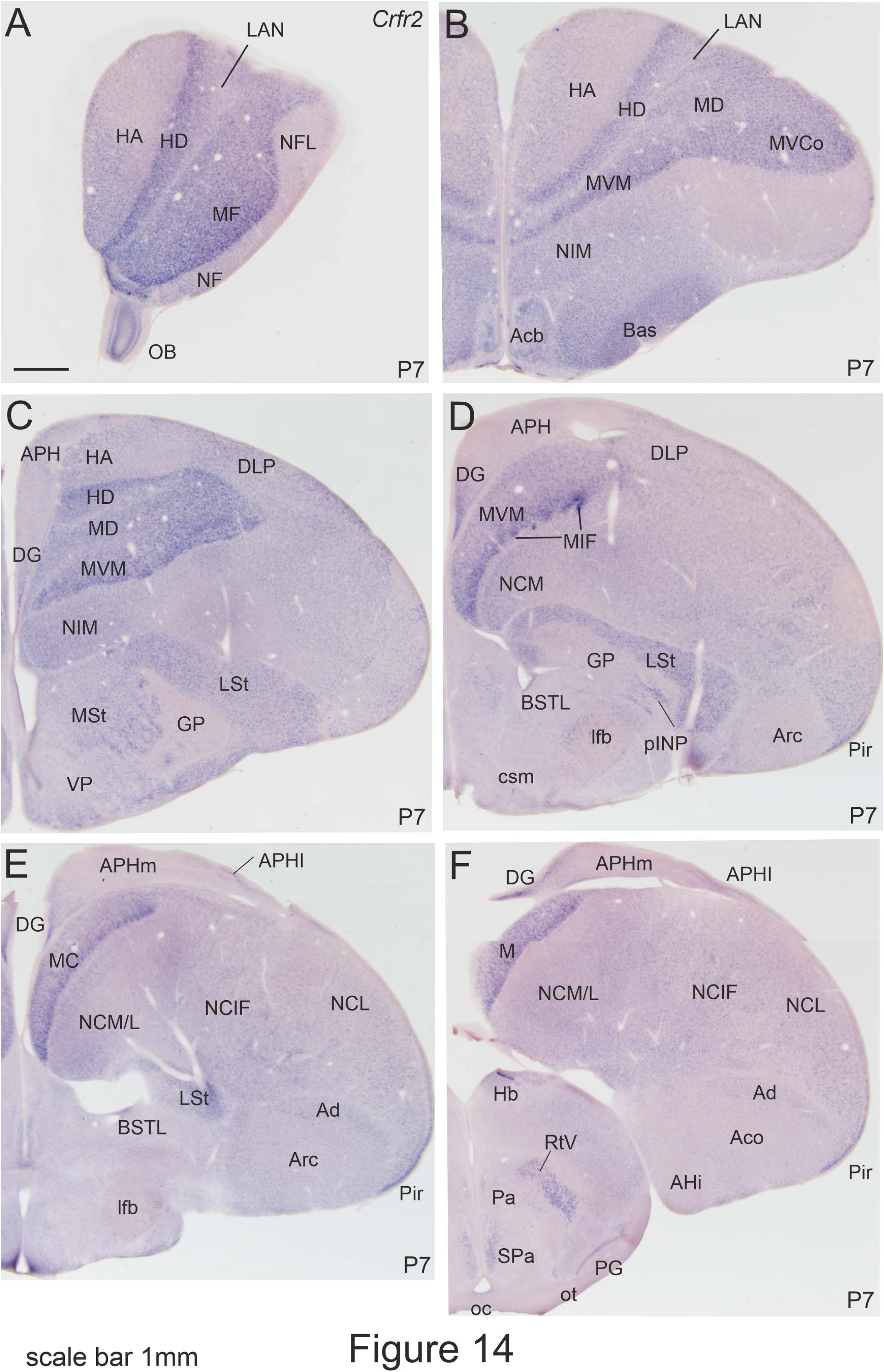
Images of frontal sections through the telencephalon and anterior hypothalamus of chicken at P7, from anterior to posterior levels, showing hybridization signal for chicken Crfr2. The expression pattern is similar to that seen at P0. See more details in text. For abbreviations see list. Scale bars: A = 1mm (applies to all).

### Chicken CRF binding protein mRNA (*cCrfbp or Crfbp*)

During embryonic and early posthatching development, the expression pattern of *Crfbp* was very different to those of both CRF receptors (Figs. 15-18). At E8, the only telencephalic region with expression was the developing striatal capsule and adjacent lateral striatum (LSt), which displayed moderate signal of the CRF binding protein mRNA (Fig. 15A,B).

**Figure 15.**
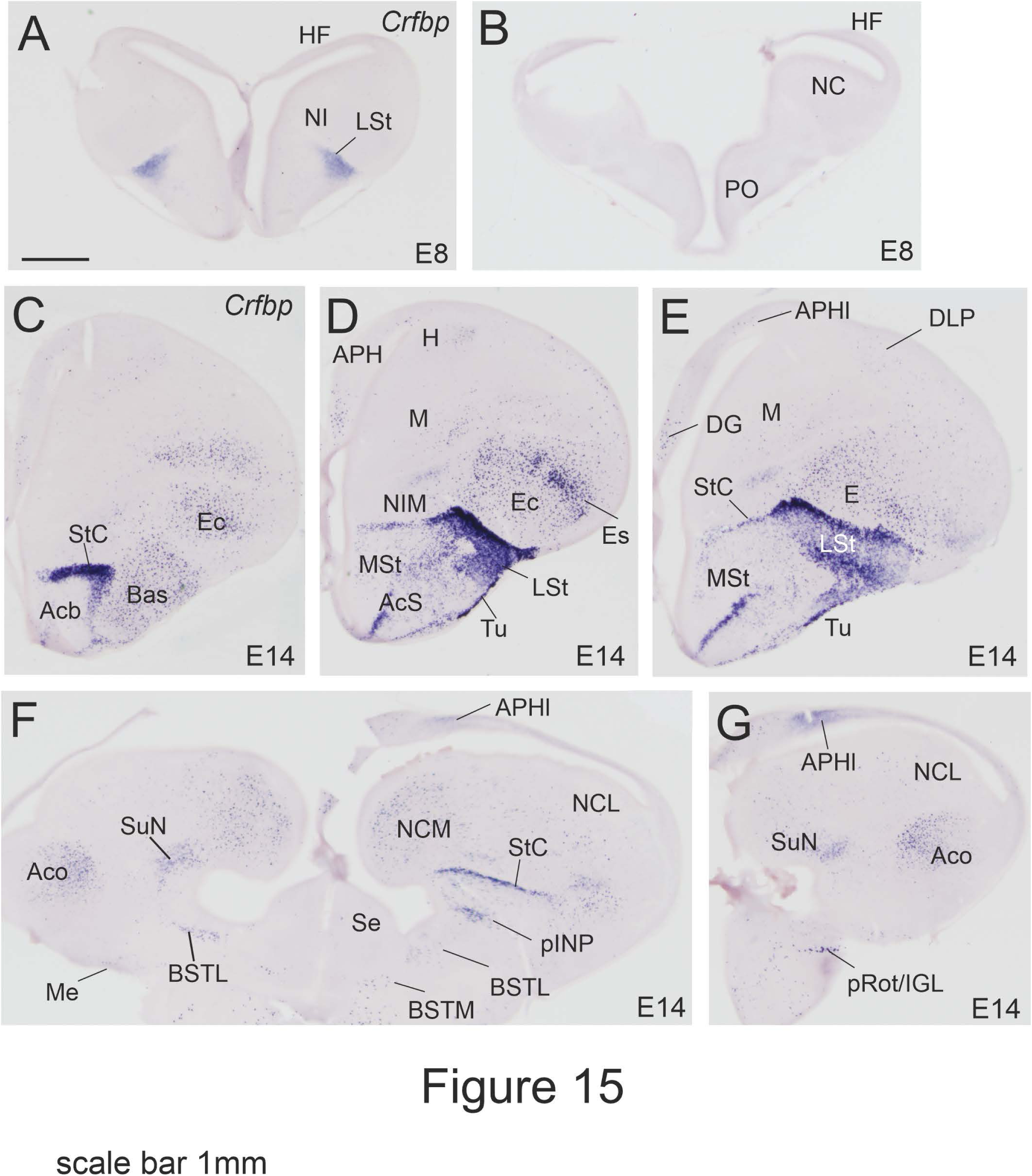
Images of frontal sections through the embryonic telencephalon and anterior hypothalamus of chicken (E8 and E14), from anterior to posterior levels, showing hybridization signal for chicken Crfrbp. Note the moderate signal in part of the striatum from E8, which becomes intense at E14, encompassing the striatal capsule and adjacent lateral striatum, as well as part of the central extended amygdala (pINP). Most of the pallidum shows light or no expression at these embryonic ages, except for the posterior levels of APHl in the hippocampal formation. See more details in text. For abbreviations see list. Scale bars: A = 1mm (applies to all).

Signal in these areas (StC and LSt) became remarkably high by E14 (Fig. 15C-E), decreasing at caudal levels (Fig. 15F). At E14, high expression was also observed in part of the accumbens shell (AcS) and in the olfactory tubercle (Tu; Fig. 15D). In addition, *Crfbp* was moderatelyexpressed in part of the peri/post-intrapeduncular island field (pINP) and light expression was observed in in part of BSTL (Fig. 15F). Minor subsets of cells were also present in the medial amygdala and BSTM (Fig. 15F). In the pallium, *Crfbp* expression was light or moderate and mostly present only in restricted parts of the medial pallium, nidopallium and arcopallium. In the nidopallium, most subpopulations of *Crfbp* expressing cells were located in its sensory nuclei including nucleus basorostralis (Bas; Fig. 15C), the entopallium and entopallial belt (E, Es; Fig. 15D,E), and field L in the caudomedial nidopallium (NCM; Fig. 15F). Expression was also observed in the subnidopallium, ventral to caudal nidopallium (SuN; Fig. 15F). In the medial pallium, minor subpopulations of *Crfbp* expressing cells were observed in the dentate gyrus (DG) and lateral parahippocampal area (APHl), and signal in the latter increased towards caudal levels (Fig. 15E-G). In the arcopallium, most of the *Crfbp* expressing cells were found in the medial core subdivision (ACoM; Fig. 15G). The core nucleus of the mesopallium (MVco) and the hyperpallium (H) also contained minor subsets of *Crfbp* expressing cells (Fig. 15C,D), but the rest of the pallium remained free of expression or contained extremely few expressing cells.

At E18 (Fig. 16A-C) and at P0 (Fig. 16D-H), the expression increased a bit in the pallium, but decreased in the supallium, while maintaining a similar pattern to that described at E14. In the hyperpallium, a moderate number of *Cfrbp* expressing cells were observed in the apical hyperpallium (HA), with a trend to be more abundant in lateral levels (Fig. 16A, D, E). In the nidopallium, in addition to the cells of the sensory nuclei (nucleus basorostralis, entopallium, field L), expression was also observed in a subset of cells of the island field of the caudal nidopallium (NCIF; Fig. 16B,G,H). In the supallium, *Crfbp* expression was similar to that at E14, but decreased in the lateral striatum, where expression became restricted to its caudal pole (Fig. 16G). In the septal region, signal was observed in the diagonal band nucleus (vertical part; DBV), around the cortico-septo-mesencephalic tract (csm), and in the pallidoseptal area (PaS; Fig. 16B,G).

**Figure 16.**
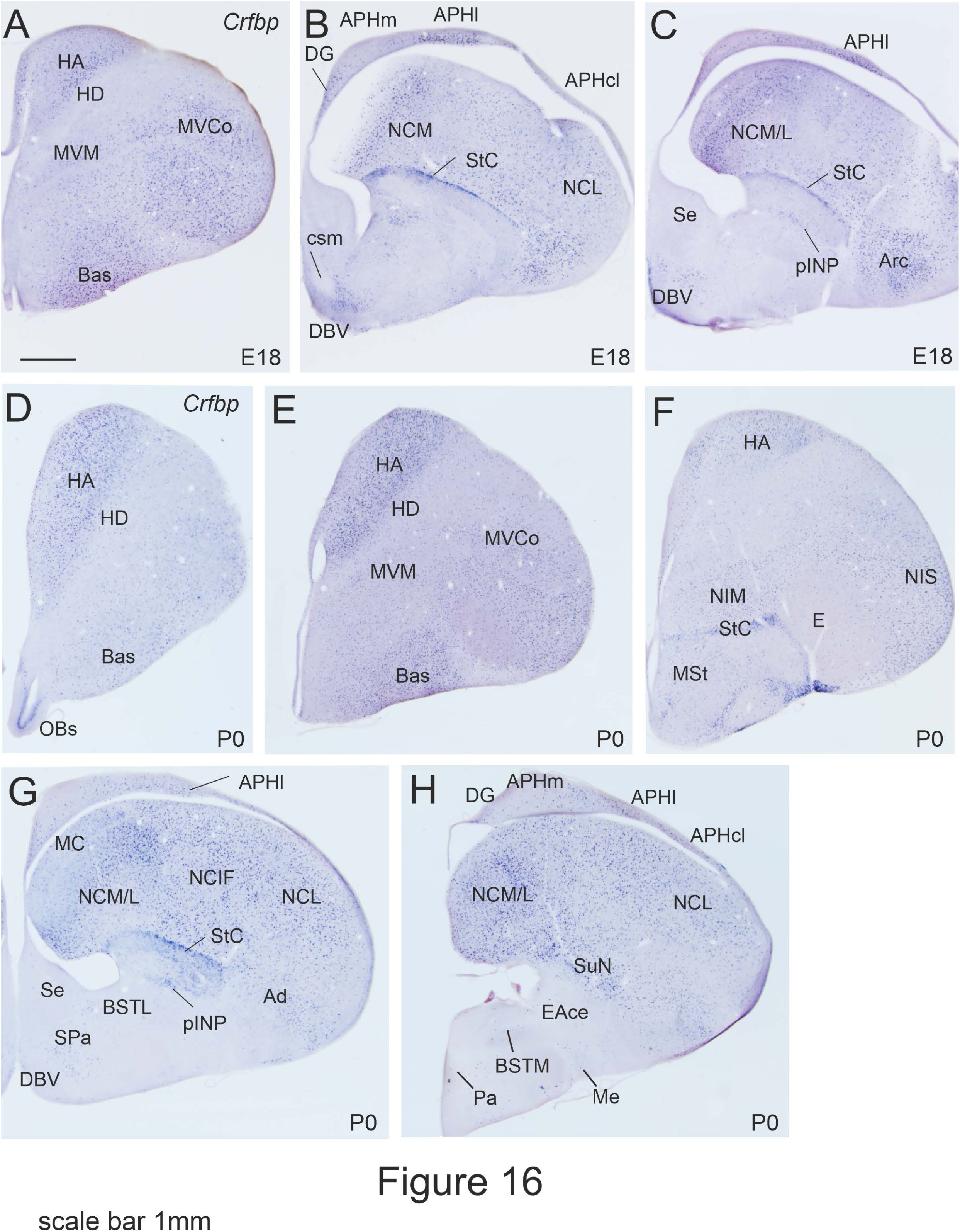
Images of frontal sections through the telencephalon and anterior hypothalamus of chicken (E18 and P0), from anterior to posterior levels, showing hybridization signal for chicken Crfrbp. Expression in the subpallium is less intense compared to E14, but is seen in similar areas. Regarding the pallium, at E18 and P0 expression is moderate in some parts of the pallium, including parts of the hippocampal formation (APHl), hyperpallium (HA), mesopallium (MVco), and nidopallium (NCM, NCL). See more details in text. For abbreviations see list. Scale bars: A = 1mm (applies to all).

By P7 (Fig. 17, details in Fig. 18), the pattern of expression remained quite similar to that at hatching, but now the expression was a bit higher in the medial pallium, arcopallium, and the basal ganglia of the subpallium. In the medial pallium, in addition to the expression in the dentate gyrus and lateral parahippocampal area, very few and scattered *Crfbp* expressing cells were seen in the medial parahippocampal area (APHm; Fig. 18A). In the arcopallium, *cCrfbp* expressing cells were seen in several subdivisions, such as the dorsal part (AD), the core nucleus (ACo), the amygdalopiriform (APir) and the amygdalohippocampal (AHi) subdivisions (Figs. 17F; 18B). In the basal ganglia, in addition to the cells found in the lateral striatum and its capsule (LST, StC), expressing cells were also seen in the medial striatum (MSt) at intermediate/caudal levels, and very few cells were seen in the pallidum (Fig. 18C,D). As before, in the extended amygdala, expressing cells were present in the peri/post-intrapeduncular island field (pINP; Figs. 17D,E; 18D). As before, a few cells were also observed in the lateral BST (Fig. 18D).

**Figure 17.**
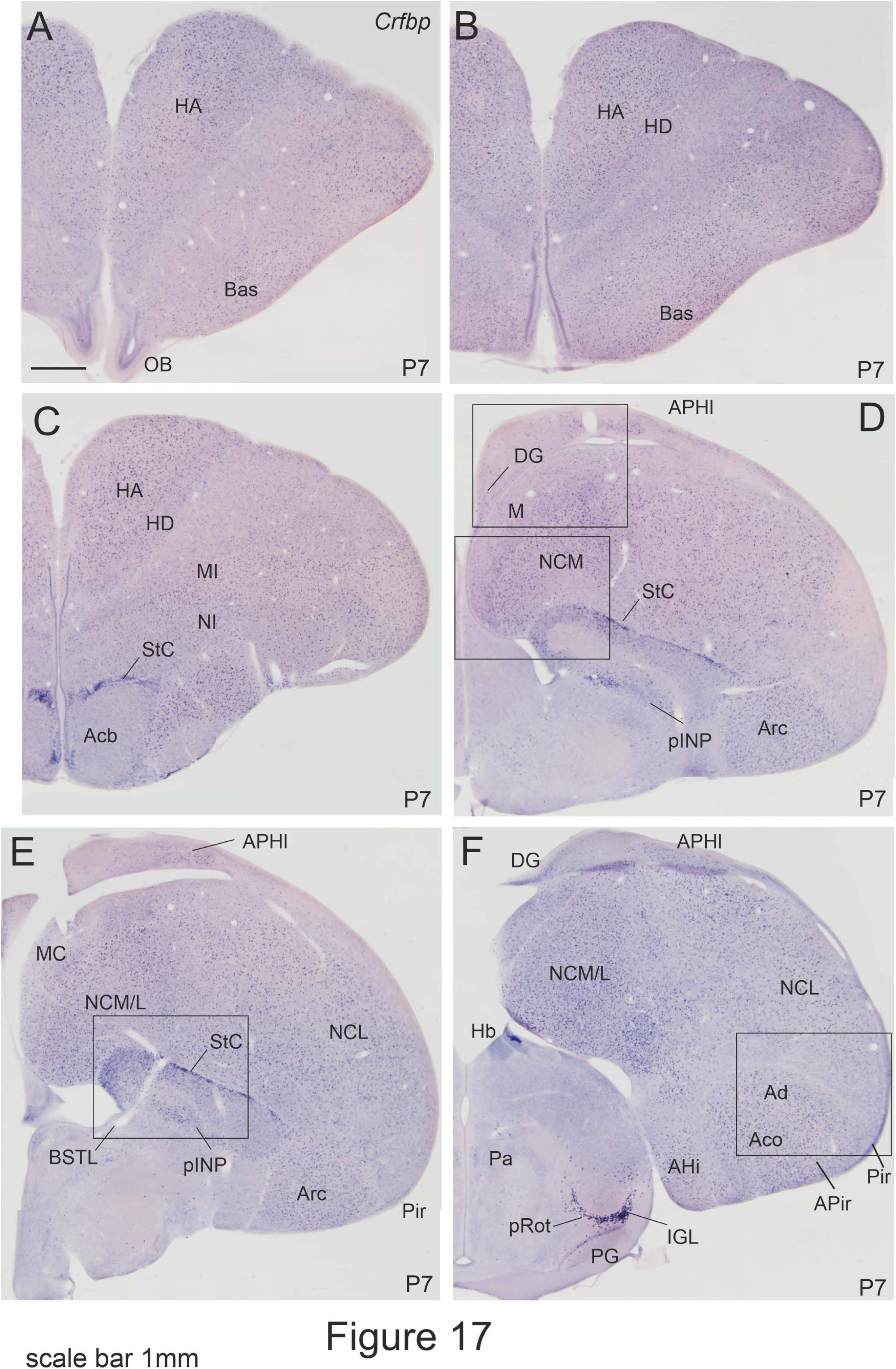
Images of frontal sections through the telencephalon and anterior hypothalamus of chicken at P7, from anterior to posterior levels, showing hybridization signal for chicken Crfrbp. Expression pattern in the subpallium is similar to that seen at hatching, while in the pallium is more spread. Details of the squared areas in D-F are shown in Figure 18. In addition to the expression in the telencephalon, high signal is observed in the thalamic perirotundic belt and intergeniculate leaflet (F). See more details in text. For abbreviations see list. Scale bars: A = 1mm (applies to all).

**Figure 18.**
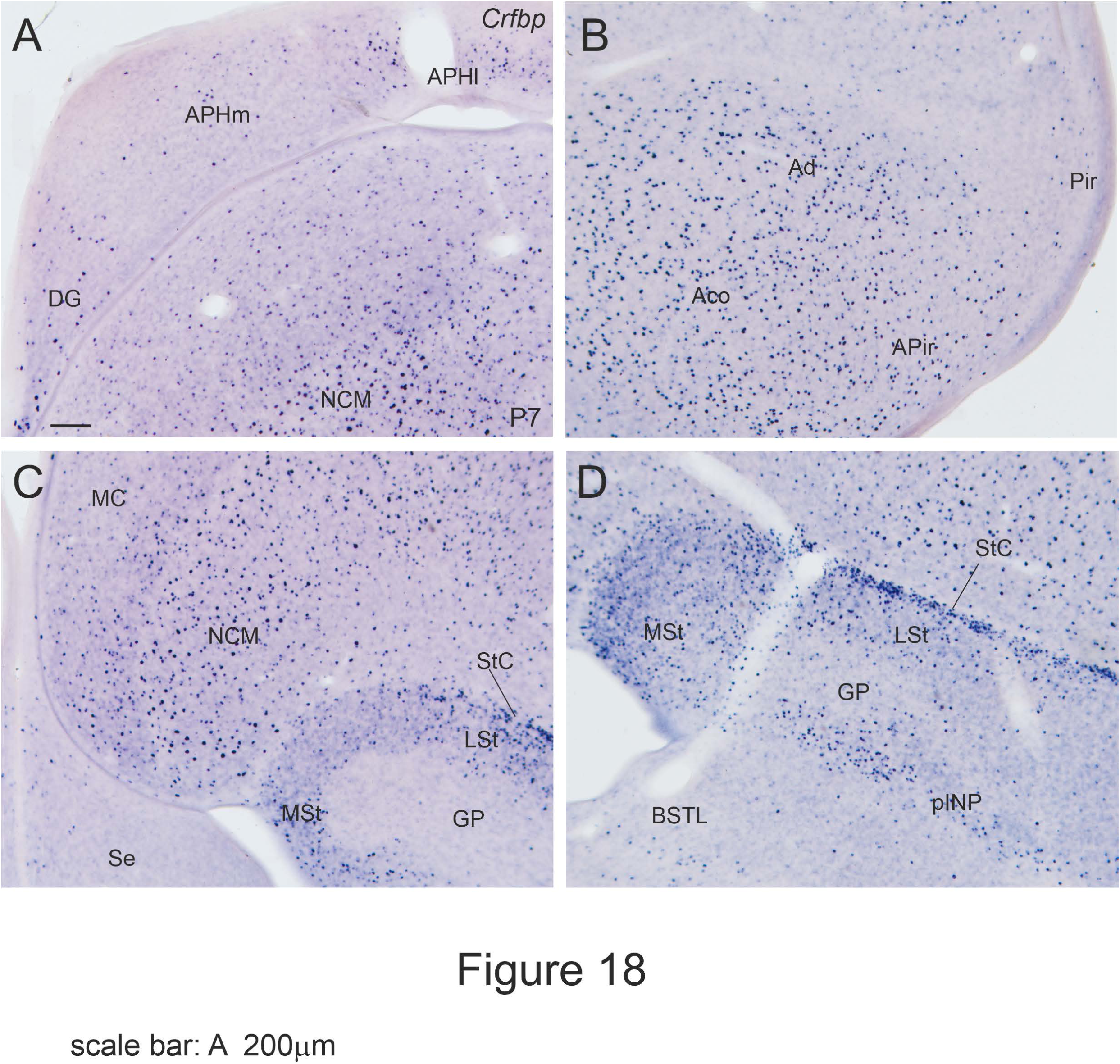
Details of the expression of Crfbp in the telencephalon of chicken at P7, from the squared areas in some sections shown in Figure 17. A shows a detail of the hippocampal formation (from the section in Fig. 17D); B shows a detail in the arcopallium (from the section in Fig. 17F); C shows a detail in the caudomedial nidopallium (from the section in Fig. 17D); D shows a detail in the subpallium (showing StC, LSt, and pINP), from the section in Fig. 17E). See more details in text. For abbreviations see list. Scale bars: A = 1mm (applies to all).

## Discussion

### 1. General findings

The major focus of our study was to map CRF systems in the chicken telencephalon during development, since CRF and its receptors are expressed in several telencephalic areas known to regulate HPA, as well as emotional and cognitive aspects of stress in other vertebrates (**Bale and Vale, 2004; Henckens et al., 2016; Herman et al., 2016**). In mammals, CRF systems of the telencephalon are involved in modulation of arousal, reward, emotion and memory formation (**Dedic et al., 2018**). This may be similar in birds, based on the expression patterns of CRF and its receptors in similar areas of the pallium and subpallium, as discussed in separate subheadings below.

Our results demonstrate mRNA expression of Crf, Crfr1, Crfr2 and Crfbp in the chicken telencephalon from early embryonic stages, at least from embryonic incubation day 8 (E8). At middle embryonic ages (E14), intense Crf expression is observed in several sensory areas of the telencephalic pallium, such as the visual areas of the hyperpallium and nidopallium (entopallium), and the multisensory and auditory areas of the nidopallium (medial intermediate nidopallium and field L, respectively). At E14, Crfr1 and Crfr2 expressions are intense in associative areas of the pallium, including areas important for sensorimotor integration and emotion processing (such as the apical hyperpallium, mesopallium, caudolateral nidopallium and arcopallium), showing partially alternating patterns. Both receptors also show moderate to intense expression in the hippocampal formation, a structure involved in memory formation and stress control. Expression of Crf receptors in the pallium continues to increase during embryonic and posthatching development, and this is accompanied by an increment in Crfbp expression. In contrast, expression of Crf and its receptors in the subpallium is generally low at middle embryonic stages, with a few exceptions (such as part of the corticopetal basal magnocellular complex), although Crfbp signal is high in parts of the subpallium from middle embryonic stages. Expression of Crf and its receptors in the central extended amygdala (a region playing a critical role in fear and anxiety) increases near hatching, and continues rising after hatching. The results point to an early development of Crf systems in pallial areas regulating sensory processing, sensorimotor integration and cognition, and a late development of Crf systems in subpallial areas regulating the stress response. However, Crfbp buffering system develops late in the pallium and early in the subpallium.

In mature mammals and birds, CRF is a major regulator of the neuroendocrine (HPA) and autonomic components of the stress response (**Vale and Bale, 2004; Carsia et al., 1986; Bons et al., 1988; Ball et al., 1989; Vandenborne et al., 2005**; **Didac et al., 2018**). This may be similar in all amniotes based on the presence of CRF immunoreactive cells in the paraventricular hypothalamic nucleus and fibers coursing to and terminating in the median eminence in different reptiles (**Mancera et al., 1991; López-Avalos et al., 1993**). In reptiles, some CRF immunoreactive fibers also reach the neurohypophysis, which may be due to coexpression with vasotocin or mesotocin in part of the CRF cells of the paraventricular hypothalamic nucleus (**Mancera et al., 1991)**, resembling the situation in mammals (**Sawchenko et al., 1984**) and birds (**Péczely and Antoni 1984**). Our results and previous studies in chicken indicate that the HPA might be active at least from middle embryonic stages, if not earlier, since CRF immunoreactive cells are present in the paraventricular hypothalamic nucleus at E14 (**Józsa et al., 1986),** and Crf and Crfr1 mRNAs are observed in this nucleus from E8 (present results). CRF immunoreactive fibers are also seen in the chicken median eminence at E14 (**Józsa et al., 1986**). Moreover, at E14 the chicken pituitary starts to influence adrenocortical activity (**Jenkins and Porter, 2004**).

In addition to activating HPA, in chicken and other non-mammals CRF has another role in food intake regulation by activating the hypothalamo-pituitary-thyroid (HPT) axis (**Bu et al., 2019**; see also **De Groef et al., 2003, 2005**). While HPA is activated by CRF binding to CRFR1 in corticotrope cells of anterior pituitary, HPT is activated by CRF2 binding to CRFR2 in thyrotrope cells of the anterior pituitary. Concurrent activation of both HPA and HPT is observed following psychogenic stress in chicken, and in this case HPT activation may be needed to generate enough energy to cope with the stressful situation (**Kadhim and Kuenzel, 2022**). In contrast to non-mammalian vertebrates (**De Groef et al., 2005, 2006**), CRF might not have such dual actions on HPA and HPT in mammals, since one study in mouse found no mRNA expression of CRFR2 and very poor mRNA expression of CRFR1 in thyrotrope-like cells of the anterior pituitary (**Westphal et al., 2009**). Nevertheless, it is possible that CRF receptor expression is induced in these cells of the anterior pituitary under specific developmental and physiological conditions (**De Groef et al., 2005, 2006**; **Westphal et al., 2009**).

Our results on the distribution of Crf mRNA expressing perikarya in different pallial and subpallial areas of the early posthatch chicken telencephalon agree with data on CRF expressing cells in early posthatch and adult chicken and quail, based on immunohistochemistry (**Richard et al., 2004**). However, we provide new details on cell groups of the pallium and subpallium, and in some of the areas we found more expressing cells than in the description by **Richard et al. (2004**): for example, in the auditory field L of the caudomedial nidopallium, we found abundant perikarya, while **Richard et al. (2004**) only described a few. This is likely due to differences in the technique employed, since we used in situ hybridization to detect mRNA (very sensitive to detect neuropeptidergic perikarya; present results), while they employed immunohistochemistry to detect the peptide (a technique than often requires pretreatment with colchicine to reach detectable levels; **Richard et al., 2004; also Ball et al., 1989**). In contrast, immunohistochemistry is better than in situ hybridization for determining the presence of the peptide/protein as well as for analyzing fiber systems (**Richard et al., 2004**). Thus, both types of approaches provide relevant, complementary information and should be considered for a better understanding of CRF systems. In general, the distribution pattern of CRF cells in the telencephalon was similar between birds and mammals (as also discussed by Richard et al., 2004), although we found a higher density of cells in some sensory pallial areas, as discussed below.

Regarding CRF receptor expression, we provide the first detailed map in the avian telencephalon. Like in mammals (**Bale and Vale, 2004; Henckens et al., 2016**), Crfr1 expression is more widespread than Crfr2 in the chicken telencephalon (present results). This suggests that these receptors are mostly segregated to different functional subsystems in mammals and birds. However, there are some differences in the expression of these receptors between both vertebrate groups. For example, in rats Crfr2 is abundant in the lateral septum, while Crfr1 is poor in this structure (**Chalmers et al., 1995**), but this is the opposite in chicken (present results). In rats, both receptors are expressed in partially overlapping areas of the olfactory system and hippocampal formation, but they segregate to different subdivisions of the pallial amygdala; moreover, the rat neocortex only expresses Crfr1 but not Crfr2 (**Chalmers et al., 1995)**. In chicken, both receptors overlap in areas of the olfactory system, hippocampal formation and the oval subnucleus of the mesopallium, but they mostly segregate to different areas in the rest of the pallium: Crfr2 is preferentially found in densocellular hyperpallium and mesopallium, while Crfr1 is abundant in apical hyperpallium, nidopallium and arcopallium. The possible function of CRF systems in these different areas is discussed below.

### Sensory, high-order association and sensorimotor integration areas

Our results agree with those of **Richard et al. (2004**) on the presence of CRF expressing cells in several first and second order sensory areas of the pallium, such as the olfactory bulb and several olfactorecipient pallial areas, the visual/somatosensory interstitial nucleus of the apical hyperpallium (IHA), the visual entopallium (mostly in its periphery) and the auditory field L. We also found CRF expressing cells in the multimodal sensory area of the medial intermediate nidopallium (NIM), which integrates visual, somatosensory and auditory signals (**Wild, 1987**; **Funke, 1989**; **Shimizu et al., 1995**; **Kröner and Güntürkün, 1999)**. One striking new finding of the present study is that abundant cells expressing Crf mRNA are observed in all of the above-mentioned areas from middle embryonic stages. In addition, our data also show high levels of Crf receptors from early embryonic stages in many of the chicken pallial sensory areas that contain Crf cells.

Receptors in thalamorecipient sensory areas of the pallium may mediate (at least partially) and/or modulate CRF-related transmission in thalamopallial projections, which agrees with the presence from E14 of high numbers of Crf expressing cells in some of the sensory thalamic nuclei that project to the pallium. Regarding the lemnothalamus, from E14 we observed CRF cells in the retinorecipient anterior dorsolateral thalamic nucleus, which is comparable to the lateral geniculate nucleus of mammals and projects to the IHA in birds (**Miceli et al., 1990**; **Shimizu et al., 1995**); and in the somatosensory dorsointermediate ventral anterior [DIVA] nucleus and somatomotor input from the ventral intermediate [VIA] nucleus, which project to the frontal pole of IHA (**Wild, 1987; Funke, 1989; Medina et al., 1997**). In the collothalamus, the shell of nucleus ovoidalis, conveying auditory information to part of field L and to caudal mesopallium (**Durand et al., 1992; Kröner and Güntürkün, 1999**; **Atoji and Wild, 2012**), also contains CRF immunoreactive cells (**Richard et al., 2004**).

Furthermore, CRF cells in these sensory pallial areas in birds may contribute to the modulation of sensory transmission and its integration within the pallium. In addition to the receptors found in first-order sensory areas, our results show the presence of abundant Crf receptors in many association and sensorimotor integration pallial areas, from early or middle embryonic stages. For example, all olfactorecipient association areas of the pallium, including the prepiriform, piriform and entorhinal cortices (**Reiner and Karten, 1985; Atoji and Wild, 2014**), contain Crfr1 and Crfr2. In the mesopallium, abundant receptors are found in the core nucleus of the ventrolateral mesopallium, MVco (also abbreviated MVL), which receives visual input from the entopallium **(Krützfeld and Wild, 2004**), and projects to the caudolateral nidopallium and arcopallium (**Atoji and Wild, 2012**). In addition, Crf and their receptors are found in areas involved in sensorimotor integration of the arcopallium (**Davies et al., 1997; Kröner and Güntürkün, 1999)** and apical hyperpallium (**Wild, 1987, 1997; Shimizu et al., 1995; Medina et al., 1997; Kröner and Güntürkün, 1999**; **Wild and Williams, 2000; Medina and Reiner, 2000**). Our results also demonstrate the presence of Crf expressing cells, as well as Crfr1 and/or Crfr2 in high-order associative pallial areas involved in different types of learning and memory, such as the caudolateral nidopallium (involved in associative learning, value coding, and working memory; **Güntürkün, 2005**; **Dykes et al., 2018**), the medial intermediate mesopallium (involved in filial imprinting, among other functions; **Horn et al., 1985**; **Horn, 1998**; **McCabe and Horn, 1994**) and the hippocampal formation (involved in spatial navigation, memory, and stress control; **Gagliardo et al., 2001; Hough and Bingman, 2004**; **Bingman et al., 2006**; **Sherry et al., 2017; Smulders, 2021**). All of these areas are connected with the arcopallium (**Kröner and Güntürkün, 1999; Csillag et al., 1994; Atoji and Wild, 2012**). The arcopallium, together with the caudal nidopallium, is considered part of the avian pallial amygdala based on embryonic origin, topological location, and transcription factors expression during development, as well as other features (reviewed by **Martínez-García et al., 2007**; **Medina et al., 2022, 2023**). Like the amygdala of mammals (**Swanson et al., 1983**; **Chalmers et al., 1995**; **Van Pett et al., 2000**; **Weera et al., 2022**), the arcopallium of chicken and other birds includes subsets of CRF expressing cells (present results; **Richard et al., 2004**) and expresses high levels of Crfr1 (present results). Moreover, it is involved in regulation of fear behavior (**Saint-Dizier et al., 2009**), passive avoidance learning (**Lowndes and Davies, 1994**; **Csillag, 1999**), taste detection (in particular, the amygdalohippocampal area, identified as nucleus taeniae in the study by **Protti-Sanchez et al., 2022**), and regulation of ingestive behavior (**da Silva et al., 2009**). In addition, it projects to the medial striatum, extended amygdala and the hypothalamus, with descending axons travelling in the ventral amygdalofugal tract (**Davies et al., 1997; Csillag, 1999; Kröner and Güntürkün, 1999; Atoji et al., 2006; Hanics et al., 2017**). In the rodent pallial amygdala (in particular, the basolateral complex), CRFR1 is expressed mostly in glutamatergic neurons, which receive input from local CRF neurons, and these receptors are critical for CRF-mediated stress responses (**Weera et al., 2022**). This may be similar in chicken and other birds. However, some differences are observed between rodents and birds, because in rodents CRF cells of the pallial amygdala appear to be interneurons (**Swanson et al., 1983**), while in birds they appear to include not only interneurons but also projection neurons. In chicken and other birds, it appears that CRF cells of the arcopallium are involved in contralateral projections through the anterior commissure and in descending projections (**Richard et al., 2004**). Interestingly, the situation in birds is similar to that described in primates (**Basset and Foote, 1992**). The consequences of this for understanding differences between species in neural regulation of stress require further investigation.

In rodents, CRF expressing cells of many areas of the pallium are interneurons, which contact pyramidal cells expressing CRF receptors (**Swanson et al., 1983**; **Chen et al., 2004a**; **Gallopin et al., 2006**). However, this is not a general rule, since the majority of the mitral cells of the olfactory bulb and their projections contain CRF (**Imaki et al., 1989**; **Basset et al., 1992**). Thus, in the pallium of mammals there are CRF neurons involved in local circuits and CRF neurons involved in extrinsic projections. A similar situation may exist in birds. Like in mammals, we found that many mitral cells of the olfactory bulb contain Crf, which may be involved in projections to several olfactorecipient pallial areas, such as the anterior olfactory area, the prepiriform/piriform cortex and the entorhinal cortex (**Reiner and Karten, 1985; Atoji and Wild, 2014**). Moreover, in chicken and quail, **Richard et al. (2004)** proposed that CRF cells of the chicken and quail arcopallium (comparable to part of the pallial amygdala) may give rise to commissural projections as well as descending projections, based on the finding of CRF immunoreactive fibers in the anterior commissure and the ventral amygdalofugal tract. Lastly, in chicken we found that Crf expressing cells in many areas of the pallium (including the IHA, the entopallium and field L) seem too abundant to only include interneurons; thus, in addition to interneurons, they might include cells involved in pallio-pallial connections, a possibility that would require further investigation.

### Auditory stimulation and synchronization of hatching

As noted above, high expression of Crf and its receptors are found in auditory telencephalic areas of chicken. What is the role of CRF in this functional system during embryonic development? In mammals, CRF is involved in acoustic startle (**Liang et al., 1992a**), and the neural circuit mediating this effect appears to involve the central amygdala, but this is not the primary center that initiates the response (**Liang et al., 1992b**). It is possible that this neural circuit also involves areas of the pallium, including the pallial amygdala, where sensorimotor integration and gating of the startle reflex in front of a sudden, intense sound takes place (**Wan and Swerdlow, 1997**). It is possible that CRF and CRF receptors in auditory-related telencephalic areas of chicken embryos contribute to modulate acoustic startle. This functional system may also mediate the long-term effects of noise-associated early life stress (**De Haas et al., 2021**). In addition, this CRF functional system may participate in modulating sound communication before hatching, and in auditory stimulation for hatching synchronization. Chicken embryos can detect auditory stimuli from E12 (**Rogers, 1995**), and appear to communicate with their mother before hatching (**Edgar et al., 2016**). Moreover, auditory stimuli from other chicks, including the clicking sound produced by egg teeth tapping during hatching, are essential for hatching synchronization (White, 1984).

### Light stimulation and lateralization

Correct light stimulation during the incubation period is known to be critical for lateralization, and conditions the side preference of pecking after hatching (**Rogers et al., 2007; Rogers, 2008**). It appears that lateralization improves cognition and the ability of performing the double task of both feed searching and predator watching, which is advantageous in natural conditions (**Rogers et al., 2004**; reviewed by De **Haas et al., 2021**). Photopigments are present in the retina from E14, and retinal photoreceptors respond to light/dark cycles from E16. However, light-sensitive cells are found from earlier stages in other brains areas, such the pineal gland and hypothalamus (**De Haas et al., 2021**). According to our results, several visual areas of the chicken telencephalic pallium show high levels of Crf expression from E12 and E14, including those in the hyperpallium and entopallium. In addition, Crf expressing cells are found from E14 in the retinorecipient anterior dorsolateral thalamic nucleus, comparable to the lateral geniculate of mammals (present results). It is possible that part of the visual pathways to the pallium and the related centers are active from middle embryonic stages, and that CRF in these areas and pathways modulates their maturation and function. CRF and its receptors may modulate the functioning of visual areas, and their prehatch expression in these and other sensory areas may contribute to their maturation, getting ready to adequately respond to sensory stimuli from hatching, which is critical in precocial animals. In addition, CRF and its receptors in these areas may mediate the effects of light-associated early life stress (**De Haas et al., 2021**).

### Corticopetal cells of the ventral telencephalon and relation to the ascending reticular activating system

One striking finding of this study is the expression of Crf mRNA in a subset of large perikarya of the so-called basal forebrain corticopetal system (located in the subpallial telencephalon). These cells were previously observed in chicken and quail using immunohistochemistry, but were proposed to belong to the extended amygdala (**Richard et al., 2004**). Like in mammals (**Alheid and Heimer, 1988**), corticopetal subpallial cells in chicken partially overlap the extended amygdala, the pallidal parts of the basal ganglia, and the lateral and medial forebrain bundles, but they belong to a different functional system characterized by projections to the pallium (**Leutgeb et al., 1996**; discussed by **Abellán and Medina, 2009**; **Kuenzel et al., 2011**). Corticopetal cells include cholinergic, GABAergic and glutamatergic neuronal subpopulations (**Medina and Reiner, 1994**; **Abellán and Medina, 2009**), and it would be interesting to investigate coexpression of these neurotransmitters with CRF. In mammals, minor subpopulations of CRF neurons in the central extended amygdala (in the sublenticular part and posterior subdivision of lateral BST) and adjacent ventral pallidum were described to coexpress glutamatergic markers (**Fudge et al., 2022**), but it is possible that these neuronal subsets -or part of them-may belong to the corticopetal system.

The presence of CRF cells in the corticopetal system is interesting because they can modulate the function of several pallial areas, and could play a role in arousal, attention and memory, functions in which mammalian corticopetal cells of the basal forebrain are involved (**Zaborszky et al., 1999**; **Saper and Fuller, 2017**). In mammals, these cells are targeted by different cell groups of the ascending reticular activating system, such as the pedunculopontine nucleus, the parabrachial nucleus, and locus coeruleus (**Zaborszky et al., 1999; Saper and Fuller, 2017**), and this appears to be similar in birds (**Kitt and Brauth, 1986**). All of these brainstem centers also contain subpopulations of CRF cells in mammals and birds (**Swanson et al., 1983; Richard et al., 2004**), so that CRF ascending systems may participate in modulating arousal, attention and memory formation in response to a stressing stimulus.

### Fear and anxiety related areas: the central extended amygdala

In mammals, two important CRF expressing neuronal subpopulations have been described in the central amygdala and related lateral part of the BST (both included in the central extended amygdala), which play important roles in fear and anxiety responses (**Koob and Thatcher-Britton, 1985**; **Nijsen et al., 2001; Daniels et al., 2004**; **Sahuque et al., 2006**; **Phelps and LeDoux, 2005**; **Ulrich-Lai and Herman, 2009; Davis et al., 2010**; **Donatti and Leite-Panissi, 2011; Dedic et al., 2018; Zhang et al., 2021**). In agreement with other studies in birds (**Ball et al., 1989; Richard et al., 2004**), our results show the presence of Crf cells in the chicken lateral BST. In addition, we found Crf cells laterally to BST, in several subdivisions that together appear to represent the avian central amygdala (as discussed by **Vicario et al., 2014, 2015, 2017; Pross et al., 2022; Medina et al., 2023**).

Like that of mammals (**Ulrich-Lai and Herman, 2009**), the lateral BST of birds is involved in fear and anxiety responses (**Nagarajan et al., 2014**). Moreover, the avian lateral BST shows connections comparable to those of mammals (**Atoji et al., 2006**), compatible with a role in regulating HPA and autonomic nervous system, and in mediating the influence on these systems of other telencephalic structures such as the hippocampal formation, the pallial amygdala and the central amygdala. Like in mammals, the lateral BST of birds is rich in GABAergic neurons, although it also contains minor subpopulations of glutamatergic cells (**Abellán and Medina, 2009**; **Vicario et al., 2015; Metwalli et al., 2022**). Depending on the neurons involved, it may lead to inhibition or disinhibition of HPA (**Ulrich-Lai and Herman, 2009**). In mammals, the lateral BST includes a subpopulation of GABAergic cells that coexpress CRF (Partridge et al., 2016), and these are particularly involved in sustained fear responses (akin to anxiety) (**Davis et al., 2010; Gafford and Ressler, 2015**). As noted above, CRF expressing cells are also found in the avian lateral BST (**Richard et al., 2004**; present results). Our results show that these CRF cells are visible from E14, suggesting that they may already be regulating HPA and possibly the autonomic system to adapt to environmental changes within the egg.

Regarding CRF cells of the central amygdala, in mammals they are involved in fear learning and memory (**Gafford and Ressler, 2015**; **Partridge et al., 2016; Asok et al., 2018**). These cells are mostly GABAergic, receive glutamatergic inputs from the pallial amygdala, and project to lateral BST, preferentially having inhibitory contacts with non-CRF cells of lateral BST, and less frequently with CRF cells of this nucleus (**Gafford and Ressler, 2015; Partridge et al., 2016**). In this study we identified similar CRF cells in the lateral parts of the avian central extended amygdala, including the peri/post-intrapeduncular island field and in the medial part of the capsular central amygdala. These cells might be comparable to those found in the mammalian central amygdala. They occupy a territory of the subpallium populated by neurons with embryonic origins and transcription factor expressions identical to those of the mammalian central amygdala, which include similar peptidergic neuronal subpopulations (**Vicario et al., 2014, 2015, 2017; Pross et al., 2022**). Moreover, this territory representing the avian central amygdala receives input from the pallial amygdala and projects to lateral BST (**Veenman et al., 1995; Kröner and Güntürkün, 1999; Atoji et al., 2016; Hanics et al., 2017**). CRF cells in the avian central amygdala might play a similar role to those of mammals in regulating the lateral BST. However, our results show that they are not visible until peri-hatching stages. Moreover, CRF receptors are not detected in lateral BST until E18 (present results). Thus, the proposed regulation of lateral BST (and indirectly of HPA) by CRF cells of the central amygdala seems to start relatively late in chicken, and this may be similar in other species. However, such regulation might occur by way of other cell types of the central extended amygdala, such as those expressing enkephalin, which are seen from early embryonic stages in chicken and zebra finch (**Vicario et al., 2014, 2015**, **2017**; **Pross et al., 2022**).

### HPA regulation: the role of the hippocampal formation

In mammals, the hippocampus exerts a negative influence on HPA by way of lateral BST (**Ulrich-Lai and Herman, 2009; Radley and Sawchenko, 2011**), and this appears to be similar in birds (reviewed by **Smulders, 2017**). Like in mammals, the avian hippocampus expresses glucocorticoid and mineralocorticoid receptors, and becomes active during stress by responding to high levels of corticosterone (reviewed by **Smulders, 2017**). The avian hippocampus also projects to the lateral BST and can thus indirectly influence HPA activity (**Atoji et al., 2006**). Since most neurons of lateral BST are GABAergic (**Abellán and Medina, 2009**), this pathway would lead to inhibition of HPA, similarly to mammals (**Ulrich and Herman, 2009; Smulders, 2017**). In mammals, a subpopulation of stress-responsive CRF interneurons is also found in the hippocampus (**Chen et al., 2004a**). These CRF expressing interneurons modulate hippocampal activation of principal cells through receptors (**Chen et al., 2004a**), and this may be similar in birds based on the presence of scattered Crf cells and abundant Crfr1 and Crfr2 in the hippocampal formation (present results). These CRF cells and both receptors are visible from E8, suggesting that CRF is modulating hippocampal function from early embryonic stages. Curiously, we found more cells during embryonic development than at hatching and at P7. One possibility is that part of the expression is transient, like in mammals (**Chen et al., 2004b**). In rats and mice, there is a transient population of Cajal-Retzius-like cells that express CRF, and during development this stress-activated neuropeptide appears to modulate dendritic differentiation of pyramidal hippocampal neurons, acting through CRFR1 (**Chen et al., 2004b**). In this context, CRF leads to decreased dendritic growth. This finding helps to explain the negative effects of CRF, if released in excess, on dendritic arbors and synaptic plasticity of hippocampus, and on memory formation (**McEwen, 1999**; **McGaugh and Roozendaal, 2002**; discussed by **Chen et al., 2004b**). This can also contribute to understanding the long-term effects of early stress in mammals and birds **(Nordgreen et al., 2006**; **De Haas et al., 2021**), as discussed in a separate section.

In birds, another telencephalic center that regulates HPA is the nucleus of the hippocampal commissure (**Nagarajan et al., 2017; Kadhim et al., 2019**). The nucleus of the hippocampal commissure contains stress-responsive CRF expressing cells (**Nagarajan et al., 2014, 2017; Kadhim et al., 2019**; present results) as well as receptors of CRF and glucocorticoids (**Kadhim et al., 2019**; present results). Compared to the control situation, this nucleus increases its activity during acute stress, but decreases its activation in chronic stress (**Nagarajan et al., 2014**). Our previous studies showed that this nucleus and the adjacent septofimbrial nucleus (both including CRF neurons from E14; present results) contain glutamatergic neurons that derive from the pallial septum, and share some similarities with the hippocampus like expression of the transcription factors Emx1 and Lhx9 during development (**Abellán et al., 2009; 2010**). Like the hippocampal formation, it is reciprocally connected with lateral and medial septal nuclei (**Atoji and Wild, 2004**). The presence of glutamatergic neurons in the nucleus of the hippocampal commissure helps to explain why its hyperactivation during acute stress leads to increased activation of HPA and increase of blood corticosterone, while its hypoactivation in chronic stress produces a decrease in both HPA activation and blood corticosterone (**Nagarajan et al., 2014**).

In mammals, corticosterone regulates the brain through glucocorticoid and mineralocorticoid receptors, such as those found in the hippocampus leading to inhibition of HPA (references above). In late posthatching and adult birds (vocal and non-vocal learners), both receptors are found not only in the hippocampus (reviewed by **Smulders, 2021**), but also in areas of the mesopallium, nidopallium, arcopallium, and subpallium, including the extended amygdala (**Matsunaga et al., 2011**; **Senft et al., 2016**). In vocal learners, expression of mineralocorticoid receptors is particularly high in telencephalic areas involved in vocal learning and control, as well as in visual and auditory nuclei of the thalamus (**Suzuki et al., 2011**; **Matsunaga et al., 2011**). In contrast, expression in generally light in the telencephalon and thalamus of non-vocal learners, although some signal is seen in visual and auditory related centers (**Matsunaga et al., 2011**). However, to our knowledge no information exists on embryonic expression in the avian brain. This information is particularly relevant in precocial species, such as quail and chicken, and is crucial to better understand regulation of the stress response as well as the effects of prehatching corticosterone (**De Haas et al., 2021**).

### Variations in stress response after hatching and relationship to imprinting

Mammals and birds pass through a hyporesponsive period to stress around birth/hatching, which coincides with an increase of corticosterone in blood; this period is followed by a rapid normalization during early posthatching (**Schapiro et al., 1962**; **Freeman and Manning, 1984**). In birds, the duration of this hyporesponsive period is about 48 hours (Freeman and Manning, 1984). Interestingly, during this period posthatchlings go through a period of filial learning or imprinting, through which they develop a social preference for parents or other individuals, and even for stimuli resembling conspecifics (reviewed by **Bolhuis, 1999**; **Rosa-Salva et al., 2015**; **Di Giorgio et al., 2017**). The brain areas and circuits involved in filial imprinting have been studied, and include one crucial area of the medial intermediate mesopallium (**Horn, 1998; Bolhuis, 1999**), as well as parts of the septum and arcopallium, including a subdivision identified as nucleus taeniae (**Mayer et al., 2017**), called amygdalohippocampal area here (as in **Puelles et al., 2019**). Interestingly, these areas express CRF receptors.

Imprinting involves high-order association and regulation of both reward and fear responses. In this regard, the medial intermediate mesopallium is a highly associative area not only involved in filial imprinting, but also in passive avoidance learning (**Rose and Csillag, 1985; Bourne et al., 1991; Dermon et al., 2002**). This area is reciprocally connected with a multisensory area of the medial intermediate nidopallium, and projects to a rostromedial accumbens-like subdivision of the striatum that is involved in motivation and reward **(Atoji and Wild, 2012**; **Bálint and Csillag, 2007**; **Bálint et al., 2011**). The medial intermediate mesopallium also projects to the arcopallium (**Csillag et al., 1994, 1997; Atoji and Wild, 2012**), a part of the avian pallial amygdala that is involved in fear responses (**Martínez-García et al., 2007; Saint-Dizier et al., 2009**). Since the arcopallium projects to the medial striatum, the medial intermediate mesopallium projection to the arcopallium also appears to be part of a loop that indirectly modulates the reward system (**Csillag et al., 1997**). All these interconnected areas contain CRF cells and/or express CRF receptors. However, while in the rostromedial striatum and arcopallium those are present from middle embryonic stages, expression of receptors in the medial intermediate nidopallium and the mesopallium starts from E18, increasing after hatching. The increment in receptor expression (in particular, Crfr1) after hatching is higher in the intermediate/caudal mesopallium. Moreover, the CRF buffering protein starts to increase in the pallium at hatching, and continues to increase moderately during the first week after hatching. It is possible that CRF transmission during the perihatching and early posthatching period contributes to shape synaptic contacts and plasticity involved in filial learning, and the higher expression levels seen in some of the pallial areas at P7 contribute to a more sophisticated modulation of the fear responses in front of strangers or other novelties.

### Early development of pallial CRF systems as a mechanism mediating long-term effects of early life stress

Several studies in mammals and birds have shown long-term negative effects of early life stress, including prenatal and early postnatal stress (**Braastad, 1998**; **Hedlund et al., 2019**; **De Haas et al., 2021**). In domestic chicken, prehatching stress is known to increase fearfulness of the offspring (**Janczak et al., 2006**), and can lead to increased injurious pecking of these animals later in life (**De Haas et al., 2021**). Moreover, injecting chicken eggs with corticosterone results in increased mortality, reduced growth, impaired laterality (**Ericksen et al., 2003**), and leads to reduced ability for filial imprinting after hatching (**Nordgreen et al., 2006**). In these cases, it appears that there is an alteration of ‘embryonic programing’, leading to changes at physiological, metabolic and epigenetic levels with long-term implications (reviewed by **De Haas et al., 2021**).

Regarding effects on brain development, to really understand the mechanisms behind short- and long-term effects of early life stress, it is mandatory to study embryonic and early posthatching expression of: 1) Glucocorticoid and mineralocorticoid receptors in brain centers known to be involved in regulation of HPA, but also in centers modulating sensorimotor integration, attention, learning and memory, motivation and emotions. 2) CRF and its receptors in the brain, since these are known to be stress-responsive and be involved in dendritic and synaptic maturation and plasticity, as well as in modulation of emotional and cognitive aspects of stress (discussion above). Considering that HPA appears to become active early in development, we can speculate that glucocorticoid/mineralocorticoid receptor expression may also develop early, at least in upper and middle centers of HPA (paraventricular hypothalamic nucleus and anterior pituitary), so that part of the negative loop regulating corticosterone production is in place from the very first moment when HPA becomes active (at about E14). Regarding development of CRF systems, our results show that there are many telencephalic centers that contain stress-responsive CRF cells and receptors from E14. In posthatching and adult animals, these centers express glucocorticoid and mineralocorticoid receptors, and it is possible that they start to express these receptors and respond to corticosterone during embryonic and/or early posthatching development.

Overall, the results help to understand the mechanisms underlying the negative effects of noise and light stress during prehatching in chicken (**Riedstra and Groothuis, 2004**; **De Haas et al., 2021**), and suggest that mechanisms for regulating stress become more sophisticated with age. We provide baseline data on the presence of CRF, receptors and binding proteins in multiple areas of the telencephalon, which open new venues for investigating their function.

## Conclusions

Corticotropin releasing factor (CRF) and their receptors play crucial roles in the regulation of physiological and endocrine responses associated to stress, which are triggered by activation of HPA and sympathetic nervous system. In mammals, several telencephalic areas, such as the extended amygdala and the hippocampus, regulate the autonomic system and the HPA. These centers include subpopulations of CRF containing neurons that, by way of CRF receptors, play modulatory roles of fear/anxiety responses and emotion-related memory formation. Our results in chicken show the presence of Crf cells and receptors in similar areas of the telencephalon, including the extended amygdala and hippocampal formation. In addition, Crf cells and/or Crf receptors are found in other parts of the pallium and subpallium involved in sensory processing and integration, arousal, reward, and feeding, among other functions. Notably, Crf and Crf receptor expression in these areas is found from early or middle embryonic stages, and may contribute to early maturation of the networks for getting ready to hatching synchronization, and to adequately respond to sensory stimuli from hatching, which is critical for fast imprinting and survival in precocial animals. In addition, they may mediate the long-term effects of early life stress.

## Author contributions

AHM processed most of the material as part of his Ph.D. research project, and AA and AP contributed in the processing. AHM photographed and prepared many study figures, and analyzed the material with help of AA and LM. AHM and LM prepared the figures for the article. AHM and LM produced the first draft of the manuscript, ED contributed to the discussion, and all authors revised it and approved it.

## List of abbreviations

Ac: Anterior commissure
Acb: Nucleus accumbens
Aco: Core or intermediate part of arcopallium
AcS: Shell of Acb
Ad: Dorsal arcopallium
AHi: Amygdalo-hippocampal area
AO or OA: Anterior olfactory area
APH: Parahippocampal area
APHcl: Caudolateral APH
APHl: Lateral APH
APHm: Medial APH
APir: Amygdalo-piriform area
Arc: Arcopallium
CCS: Caudocentral septal nucleus
CoS: Commissural septal nucleus
csm: Cortico-septo-mesencephalic tract
DG: Dentate gyrus
Bas: Nucleus basorostrallis
BasSh: Shell of Bas
BMC: Basal magnocellular corticopetal complex
BST: Bed nucleus of the stria terminalis
BSTL: Lateral BST
BSTM: Medial BST
BSTM1: Dorsal subdivisión of BSTM
CeC: Capsular central amygdala
Ceov: Oval central amygdala nucleus
DB: Diagonal band nuclei
DLA: Anterior dorsolateral thalamic nucleus
DLI: Anterior dorsointermediate thalamic nucleus
DLM: Anterior dorsomedial thalamis nucleus
DLP: Dorsolateral pallium
E: Entopallium
EA: Extended amygdala
EAce: Central EA
Ec: Entopallial core
Ent: Entorhinal cortex
Es: Entopallial shell or belt
GP: Globus pallidus
Gr: Granular cell layer of OB
H: Hyperpallium
HA: Apical hyperpallium
Hb: Habenula
HD: Densocellular hyperpallium
HF: Hippocampal formation
HiC: Nucleus of the hippocampal commissure
IGL: Intergeniculate leaflet
IHA: Interstitial nucleus of HA
INP: Intrapeduncular nucleus
L: Auditory field L (in NCM)
lac: Lateral limb of ac
LAN: Laminar pallial nucleus (lateral to HD and dorsal to MD; it extends into a superficial area, often referred as superficial or lateral hyperpallium)
lfb: Lateral forebrain bundle
LOT: Nucleus of the lateral olfactory tract
LSt: Lateral Striatum
M: Mesopallium
MC: Caudal mesopallium
MD: Dorsal mesopallium
Me: Medial amygdala
MF: Frontal mesopallium
Mi: Mitral cell layer of OB
MI: Intermediate mesopallium
MIF: Mesopallial island field
MIV or MV: Ventral MI
MPA: Medial preoptic area
MPO: Medial preoptic nucleus
MSt: Medial striatum
MVCo: Core nucleus of MIV
MVM: Medial MIV
N: Nidopallium
NC: Caudal nidopallium
NCIF: Island field of NC
NCL: Lateral NC
NCM: Medial NC
NF: Frontal nidopallium
NFL: Lateral NF
NI: Intermediate nidopallium
NIM: Medial NI
NIS: Superficial NI
OA: Anterior olfactory area
OB: Olfactory bulb
Pa: Paraventricular hypothalamic nucleus
PG: Pregeniculate prethalamic nucleus (also known as ventral lateral geniculate nucleus)
pINP: Peri/post-INP island field
Pir: Piriform cortex (olfactory cortex)
PO: Preoptic region
PPir: Prepiriform cortex
pRot: Perirotundic belt or area
RB: Retrobulbar area
RtV: Reticular prethalamic nucleus, ventral part
Se: Septum
SFi: Septofimbrial nucleus
SLI (or LSI): Intermediate subdivision of lateral septum
SLV (or LSV): Ventral subdivision of lateral septum
SO: Supraoptic nucleus
SPa: Septopallidal transition area
StC: Striatal capsule
SuPO: Subpreoptic area (part of the telencephalonopto-hypothalamic domain or TOH)
Th: Thalamus
vaf: ventral amygdalofugal tract
VDB: Vertical subnucleus of DB
VP: Ventral pallidum

## Acknowledgments

We deeply thank all Agencies that funded our research. We also thank the technicians and other staff of the Department of Experimental Medicine.

